# Immunoinformatics-Based Design of a Chimeric Vaccine Against *Necator americanus* Targeting the *Na*-APR-1, *Na*-GST-1, and *Na*-ASP-2 Vaccine Candidates

**DOI:** 10.64898/2026.10.09.757581

**Authors:** Yanick Aqua Stong Tangan, G Takop Nchanji, Kwi Pilate Nkineh, Teke Mary Efeti, Joan Amban Chick, Eric Ndumu, Lahngong Methodius Shinyuy, Marcel Nyuylam Moyeh, Stephen Mbigha Ghogomu, Robert Adamu Shey

**Affiliations:** Department of Biochemistry and Molecular Biology, Faculty of Science, University of Buea, Buea, Cameroon; Molecular and Cell Biology Laboratory, Biotechnology Unit, University of Buea, Cameroon; Tropical Disease Interventions, Diagnostics, Vaccines, and Therapeutics (TroDDIVaT) Initiative, Frontline Association, Tiko, Cameroon; VivoSpera Clinical Research Organisation, Kampala, Uganda; Department of Gerontology, Faculty of Medicine and Pharmacy, Vrije Universiteit Brussel, Laarbeeklaan 103, B-1090 Brussels, Belgium; MRC Unit the Gambia at London School of Hygiene and Tropical Medicine, Fajara, The Gambia; Centre for Research in Infectious Diseases, Yaounde, Cameroon; Pharmacognosy Laboratory, Center for Interdisciplinary Research on Medicines (CIRM), University of Liège, 4000 Liège, Belgium; Department of Computer and Information Sciences, Covenant University, Ota, Nigeria; Covenant Applied Informatics and Communication Africa Centre of Excellence (CApIC-ACE), Covenant University, Ota, Nigeria; Covenant University Bioinformatics Research (CUBRe), Covenant University, Ota, Nigeria; Centre for Malaria Elimination, Institute of Tropical Medicine, Mount Kenya University, Thika, Kenya; Department of Chemical and Biological Engineering, University of Bamenda, Bamenda, North West Region, Cameroon

**Keywords:** Hookworm vaccine, *Necator americanus*, Chimeric vaccine, Immunoinformatics, Neglected tropical disease, Multi-epitope vaccine

## Abstract

Hookworm infection, caused predominantly by *Necator americanus*, is a soil-transmitted helminthiasis that disproportionately affects impoverished communities across tropical and subtropical regions, with an estimated 406-480 million people infected globally. Control continues to rely largely on mass drug administration, an approach challenged by rapid reinfection, the absence of durable protective immunity, and growing concern over reduced drug efficacy. While the leading recombinant protein antigens *Na*-APR-1, *Na*-GST-1, and *Na*-ASP-2 have demonstrated clinical safety and immunogenicity, the withdrawal of *Na*-ASP-2 after triggering allergic reactions in individuals with prior natural exposure revealed both the promise and the risk of single-antigen approaches, underscoring the need for a rationally designed multi-antigen vaccine. We used an immunoinformatics approach to design a novel chimeric vaccine candidate by combining carefully selected B-cell and T-cell epitopes from the major clinical-stage hookworm protein antigens with an immunostimulatory adjuvant sequence. The candidate’s 3D structure was predicted, refined, and validated using established bioinformatics approaches, and its interaction with the innate immune receptor TLR4 was assessed through molecular docking and normal-mode analyses. The selected antigens showed conservation across related hookworm species, suggesting the potential for broader cross-protection. The designed candidate was predicted to be antigenic, non-allergenic, and non-toxic, and immune simulation analyses indicated its capacity to elicit coordinated humoral and cellular immune responses. HLA coverage analysis indicated broad population applicability. Docking and normal-mode analyses of its interaction with TLR4 indicated stable and favourable binding. This study offers a rational, safety-driven approach to multi-epitope vaccine design against hookworm infection, integrating the strengths of clinically evaluated protein antigens while addressing the safety concerns raised by prior single-antigen candidates. Further experimental validation is required to confirm the safety and efficacy of the vaccine construct *in vitro* and *in vivo*. These findings lay the groundwork for future efforts toward an effective immunoprophylactic strategy against this neglected disease.

## Introduction

Hookworms are soil-transmitted nematode parasites that infect humans through contact with contaminated soil. They predominantly affect impoverished populations in tropical and subtropical regions [1]. Hookworms establish chronic intestinal infections and feed on host blood, often causing iron-deficiency anemia. During larval migration, they may also induce pulmonary manifestations [2]. Globally, hookworm infection affects hundreds of millions of people, with estimates ranging from approximately 406 to 480 million infected individuals, while the Global Burden of Disease 2021 study reported 642.72 million prevalent cases [3]. Hookworm infections account for an estimated 1.38-4 million disability-adjusted life years (DALYs) lost annually. Human hookworm infections are primarily caused by two species: *Ancylostoma duodenale* and *Necator americanus* [4]. Among the causative species, *N. americanus* is responsible for approximately 79% of human hookworm infections and is the predominant species worldwide [5]. Control of the infection currently relies on mass drug administration (MDA) with benzimidazole anthelmintics; however, its long-term effectiveness is limited by rapid reinfection, the absence of durable immunity, and the emergence of drug resistance [6]. These limitations highlight the urgent need for a prophylactic vaccine, as emphasized in the WHO 2021-2030 NTD Roadmap [7].

Vaccine development efforts have largely focused on three antigens that play key roles in hookworm biology and host immune recognition: *N. americanus* Aspartic Proteinase-1 (*Na*-APR-1), *N. americanus* Glutathione S-Transferase-1 (*Na*-GST-1), and *N. americanus Ancylostoma* secreted protein-2 (*Na*-ASP-2) [8–10]. *Na*-APR-1 and *Na*-GST-1 are critical enzymes involved in hemoglobin digestion and heme detoxification, making them promising protective vaccine targets [11, 12]. Phase I clinical trials demonstrated that both candidates are safe, well tolerated, and immunogenic in hookworm-naïve and endemic populations [9, 10]. On the other hand, *Na*-ASP-2, a larval antigen secreted upon host skin penetration, experienced a clinical setback that has fundamentally reshaped hookworm vaccine strategy. Although it demonstrated safety in hookworm-naïve adults, *Na*-ASP-2 induced IgE-mediated allergic reactions in hookworm-exposed individuals, limiting its clinical development [13]. Seroepidemiological studies show that individuals, including young children in endemic areas, possess pre-existing *Na*-ASP-2-specific IgE, limiting the safe use of the intact antigen [13]. Clinical development was discontinued, underscoring the need to identify and mitigate the allergenic potential of helminth vaccine antigens before clinical application [14].

These findings support the development of multi-epitope vaccines (MEVs), which are rationally designed chimeric constructs comprising selected B-cell, cytotoxic T-lymphocyte (CTL), and helper T-lymphocyte (HTL) epitopes from multiple antigens, linked by molecular linkers and combined with an immunostimulatory adjuvant [15]. In addition, unlike whole-protein vaccines, MEV design enables the computational selection of antigenic, safe epitopes while excluding potentially allergenic or tolerogenic sequences [16]. MEV design integrates immunoinformatics tools for epitope prediction, structural analysis, docking, and immune simulation, with successful applications across various pathogens, including *Mycobacterium tuberculosis* [17, 18], SARS-CoV-2 [19], *Helicobacter pylori* [20], *Onchocerca volvulus* [21] and *Strongyloides stercoralis* [22]. Our group previously applied this immunoinformatics pipeline (unpublished data) to characterize *Na*-APR-1, *Na*-GST-1, and *Na*-ASP-2 vaccine candidates, accurately predicting *Na*-ASP-2’s allergenic risk while supporting the safety and immunogenic potential of *Na*-APR-1 and *Na*-GST-1. These findings validated immunoinformatics as a useful preclinical screening approach. They highlighted the need for a vaccine candidate that combines broad B- and T-cell epitope coverage with optimal safety and balanced immune activation.

The study aims to develop a novel chimeric vaccine candidate (designated EpiNecaVac) incorporating safe, and antigenic epitopes from *Na*-APR-1, Na-GST-1, and *Na*-ASP-2. *Na*-ASP-2 epitopes were selectively incorporated through computational screening to retain immunogenic regions while excluding allergenic and toxic sequences, potentially harnessing its vaccine potential without its clinical risks [14, 23]. The chimeric vaccine also incorporates a *Mycobacterium tuberculosis* RpfE TLR4 agonist adjuvant and linker sequences, followed by comprehensive evaluation of its safety, structural properties, immune potential, and population coverage. Overall, this study seeks to provide a rigorous, computationally validated multi-epitope vaccine candidate that leverages the immunogenic potential of all three clinical hookworm antigens while applying the safety engineering advantages of immunoinformatics to generate a safer, broader, and more immunologically potent architecture than the current individual candidates in clinical development.

## Materials and methods

### Protein selection for vaccine design

The amino acid sequences of three immunogenic proteins associated with *N. americanus* pathogenesis or virulence (accession numbers: Q9N9H3, D3U1A5, and Q7Z1H1) were retrieved from the [UniProtKB](https://www.uniprot.org/) database for preliminary analysis, epitope prediction, and chimeric antigen design [9, 10, 13]. The 3D structure of the human TLR4 (PDB ID: 4G8A) was obtained from the [Protein Data Bank](https://www.rcsb.org/) database. A 50S ribosomal protein L7/L12 (Locus RL7_MYCTU, UniProt ID: P9WHE3) was selected as a built-in adjuvant (since it is reported to be a TLR4 agonist) [24].

### Signal peptide, sequence localization and transmembrane domain prediction

The antigens were subsequently evaluated through a set of preliminary computational analyses, including signal peptide prediction with the SignalP-6.0 and [TOPCONS](https://topcons.cbr.su.se/) servers to distinguish classical secretory proteins from non-secretory proteins. Subcellular localization was predicted using the [DeepLoc 2.1](https://services.healthtech.dtu.dk/services/DeepLoc-2.1/) and [WoLF-PSORT](https://wolfpsort.hgc.jp/) servers, while transmembrane domains were predicted using the [DeepTMHMM 1.0](https://services.healthtech.dtu.dk/services/DeepTMHMM-1.0/) and [TOPCONS](https://topcons.cbr.su.se/pred/) servers.

### Analysis of protein conservation

To assess the potential of the selected proteins to provide cross-protection against other closely related hookworm species, BLAST analysis was performed against the [UniProtKB](https://www.uniprot.org/blast) database, and the percentage sequence identities of homologous proteins were determined. This rationale is based on the fact that high sequence conservation within the *Necator genus* may indicate the potential for cross-protective immunity [25]. UniProtKB is a comprehensive protein database that integrates information from diverse sources, serving as the primary repository for protein sequences and their functional annotations [26].

### Prediction of linear B-cell epitopes

B cells are key mediators of humoral immunity, and *N. americanus*-specific B-cell and antibody responses have been implicated in the host immune response and potential protection against hookworm infection [27]. Linear B-cell epitopes were assessed using the [ABCpred](https://webs.iiitd.edu.in/raghava/abcpred/) server. ABCpred utilizes recurrent neural networks to assess linear B-cell epitopes with high specificity, and for this study, a strict cutoff (≥0.70) was used to ensure precision [28]. To ensure high-confidence epitope identification, only peptides with prediction scores ≥ 0.70 were selected for subsequent analyses.

### Prediction of cytotoxic T lymphocytes (CTL) epitopes

Cytotoxic T-lymphocyte (CTL) epitopes were predicted using the [NetCTL 1.2](https://services.healthtech.dtu.dk/services/NetCTL-1.2/) server, which integrates three key parameters involved in antigen presentation: MHC class I peptide binding affinity, proteasomal C-terminal cleavage, and transporter-associated antigen processing (TAP) transport efficiency, to identify potential CTL epitopes [29]. Though the server supports predictions across 12 MHC class I supertypes, only the A2, A3, and B7 supertypes were selected for this study since together they provide up to 90% population coverage [30]. Artificial neural network-based models are employed to predict MHC class I binding and proteasomal cleavage, whereas TAP transport efficiency was assessed using a weight matrix approac [31]. A default threshold score of 0.75 was used for the identification of CTL epitopes. CTL epitopes were included based on emerging evidence that CD8⁺ T cells contribute to protective immunity against helminths primarily through cytokine production rather than direct cytotoxic activity [29].

### Prediction of Helper T-cell (HTL) epitopes

Helper T-cell epitopes were assessed using the NetMHCII-2.3 server. 15-mer peptides were evaluated for their binding affinity to human HLA-DR, HLA-DQ, and HLA-DP alleles using the server’s artificial neural network-based prediction algorithm [32]. Default prediction parameters were applied, and epitopes were sorted as strong binders (SB) or weak binders (WB) [33]. HTL epitopes binding to HLA-DR, HLA-DQ, and HLA-DP alleles were assessed for all selected antigens. Only highly promiscuous epitopes, defined as those capable of binding multiple HLA class II alleles and classified as strong binders (SB), were selected for inclusion in the multi-epitope vaccine construct.

### Prediction of epitope antigenicity

In order to ensure that the designed chimeric vaccine candidate can elicit a robust immune response, the [VaxiJen v2.0](https://www.ddg-pharmfac.net/vaxijen/VaxiJen/VaxiJen.html) server was employed to predict the antigenicity scores of all potential epitopes. VaxiJen employs an alignment-independent approach that predicts antigenicity based on the physicochemical properties of peptides and proteins. It was selected for this study because its auto-cross-covariance (ACC)-based transformation enables reliable antigenicity prediction, even for novel or synthetic sequences [34]. A default threshold of 0.6 was set for the parasite model and was used to assess epitope antigenicity, and only those with scores above 0.6 were chosen to construct the chimeric antigen. **Figure 1** shows the various steps involved in this study.

**Figure 1.**
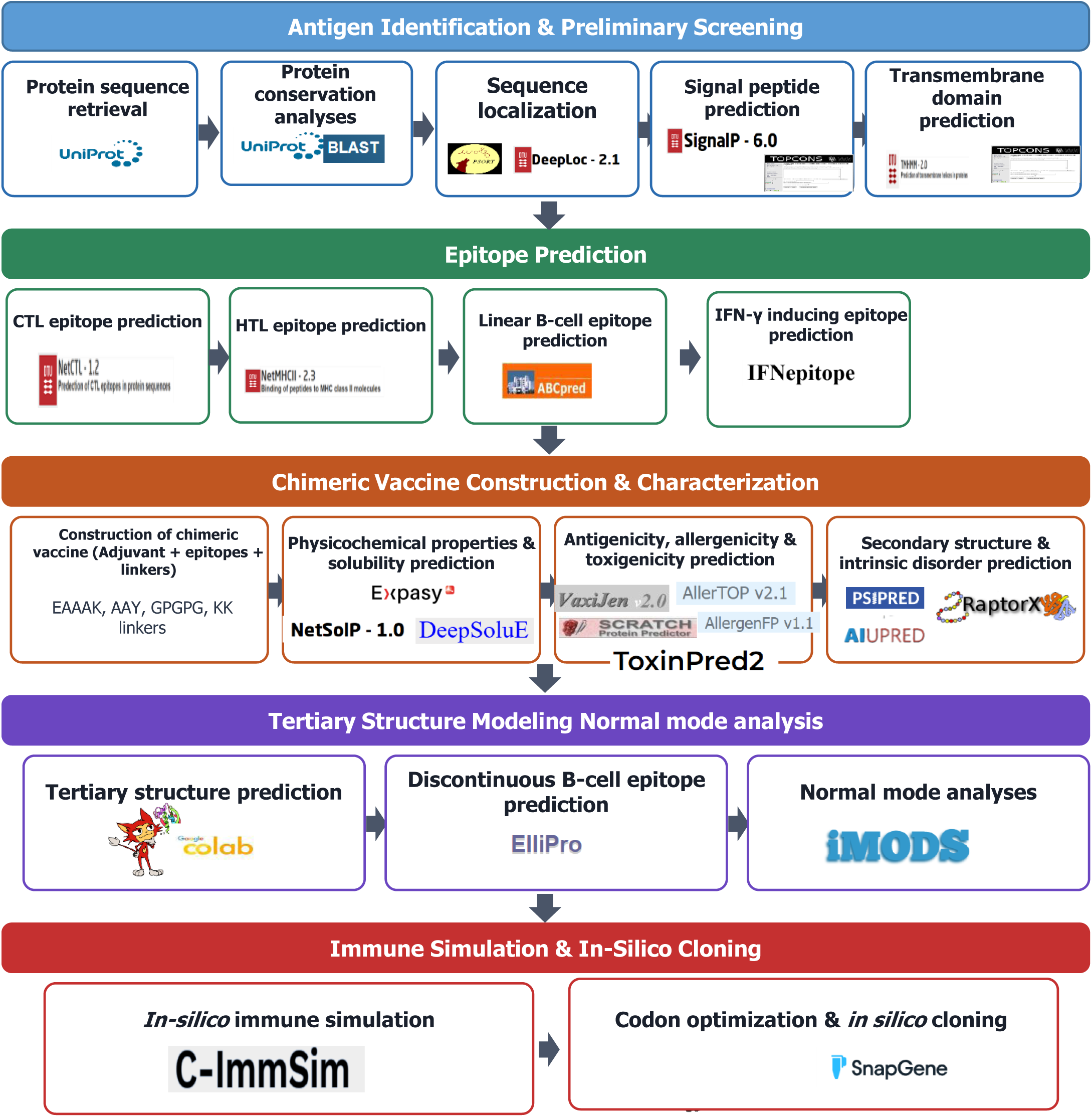
A flowchart for the designed study showing the key steps. The entire approach used in the study comprises several phases, which began with the identification and retrieval of the target protein antigens and conservation analysis. Following the preliminary screening for localization, signal peptide, and transmembrane regions, epitope predictions (CTL, HTL, and linear-B cell epitopes) from selected proteins were performed. The vaccine candidate was designed, followed by 3D modeling, refinement, and validation. Molecular docking of the designed vaccine candidate and TLR4, was followed by normal mode analysis. *In silico* immune simulation to investigate the potential of the vaccine candidate to elicit an immune response was performed. Lastly, reverse translation, codon optimization, and *in silico* cloning in a suitable plasmid for bacterial expression were performed.

### Population coverage analysis

To assess the potential global applicability of the vaccine candidates, a population coverage analysis was performed using the [IEDB Population Coverage](https://tools.iedb.org/population/) tool. The IEDB Population Coverage tool assess the proportion of a population capable of mounting an immune response based on HLA allele frequencies. It supports analyses of both MHC class I- and class II-restricted T-cell epitopes and reports combined or separate population coverage, together with the average number of epitope hits and the minimum number of epitope-HLA combinations recognized [35]. For this study, MHC class I analysis was limited to the A2, A3, and B7 supertypes, while MHC class II analysis focused on the HLA-DR, HLA-DQ, and HLA-DP alleles, as these collectively provide optimal population coverage in regions endemic for necatoriasis [30, 33].

### Construction of chimeric vaccine candidate

Candidate epitopes identified from the three *N. americanus* antigens were subjected to sequential screening for antigenicity, allergenicity, and toxicity. Epitopes meeting these criteria were subsequently prioritized based on their predicted CTL, HTL, and linear B-cell properties and selected for incorporation into the chimeric vaccine construct. The epitopes were linked using AAY, GPGPG, and KK linkers for CTL, HTL, and B-cell epitopes, respectively. These linkers facilitate appropriate epitope separation, minimizing the formation of junctional epitopes while preserving the immunogenicity, structural integrity, and accessibility of individual epitopes to the immune system [36]. Moreover, the 50-S ribosomal protein L7/L12 (Locus RL7_MYCTU, UniProt ID: P9WHE3), reported to be a TLR4 agonist [24], was incorporated as a built-in adjuvant) to enhance the vaccine’s immunogenicity and was added at the N-terminus through an EAAAK linker. TLR4 plays a crucial role in mediating protective immune responses against helminth infections, including hookworm disease [37, 38]. The sequence of the adjuvant was obtained from the [UniProtKB](https://www.uniprot.org/) database. The EAAAK linker has been reported to enhance protein expression, stability, and biological activity [39]. GPGPG linkers have been widely used in epitope-based vaccine constructs to reduce the formation of junctional epitopes, thereby facilitating efficient antigen processing and presentation [40]. AAY linkers were used to connect CTL epitopes, facilitating efficient proteasomal cleavage and promoting proper epitope separation while minimizing junctional immunogenicity and enhancing the overall immunogenicity of the vaccine construct [41]. For the linear B-epitopes, KK linkers were used. They have been reported to maintain the independent immune reactivity of each epitope [42]. In summary, all the incorporated linkers are crucial for ensuring an extended conformation (flexibility), aiding in folding, separating protein domains, and generally enhancing the stability of the recombinant multi-epitope vaccine structure [43]. Finally, a 6xHis-tag was added at the C-terminus of the chimeric vaccine candidate for application in identification and purification procedures [23].

### Prediction of IFN-γ-inducing epitopes

Interferon-gamma (IFN-γ) plays a critical role in both innate and adaptive immunity and has been identified as an important marker of Th1-mediated immune responses associated with protection against *N. americanus* infection [44]. [IFNepitope](https://webs.iiitd.edu.in/raghava/ifnepitope/predict.php) server was used to predict 15-mer IFN-γ-inducing epitopes for the designed vaccine construct. The prediction was performed by a motif and support vector machine (SVM) hybrid approach. The server utilizes a dataset comprising IFN-γ-inducing and non-inducing MHC class II-binding peptides to predict potential epitopes capable of activating T-helper cells [45].

### Prediction of epitopes for mouse MHC II alleles

To evaluate the ability of the designed vaccine candidate to induce immune responses in preclinical studies, 15-mer MHC II binding peptides for mouse alleles in the designed vaccine candidate were predicted using the [NetMHCII 2.3](https://services.healthtech.dtu.dk/services/NetMHCII-2.3/) server. The functional algorithm of this server has been described above.

### Analysis of physicochemical properties and solubility

The different physicochemical properties of the chimeric vaccine candidate were predicted using the [ExPASy ProtParam](https://web.expasy.org/protparam/) web server [46]. The tool predicts different physicochemical parameters including amino acid composition, theoretical isoelectric point (pI), molecular weight, instability index, aliphatic index, grand average of hydropathicity (GRAVY), and the predicted in vitro and in vivo half-lives following expression in *Escherichia coli*, yeast, and mammalian cells. Also, the solubility of the chimeric vaccine candidate was predicted using the NetSolP-1.0 and DeepSoluE servers. NetSolP-1.0 evaluate protein solubility and purification suitability in *E. coli* directly from sequence using deep-learning protein language models, thereby providing high predictive performance [47]. DeepSoluE tool employs a long-short-term memory (LSTM) network with hybrid physicochemical and amino acid features to predict bacterial protein solubility [48].

### Prediction of antigenicity, allergenicity, and toxigenicity of the chimeric vaccine

The antigenicity of the designed chimeric vaccine candidate was evaluated using both the [ANTIGENpro](https://scratch.proteomics.ics.uci.edu/) and [VaxiJen v2.0](https://www.ddg-pharmfac.net/vaxijen/VaxiJen/VaxiJen.html) (described above) servers. ANTIGENpro server is an alignment-free, sequence-based prediction tool trained on a large, non-redundant dataset derived primarily from protein microarray experiments [49]. In addition, the allergenicity of the designed chimeric vaccine candidate was evaluated using the AllerTOP v2.0 and AllergenFP web servers. [AllerTOP v2.0](https://www.ddg-pharmfac.net/allertop_test/) employs amino acid E-descriptors, together with auto- and cross-covariance (ACC), and k-nearest neighbors (kNN) classification to assess allergens, achieving 85.3% accuracy by 5-fold cross-validation [50]. On the other hand, the [AllergenFP](https://www.ddg-pharmfac.net/AllergenFP) server utilizes an alignment-free, descriptor-based fingerprint approach to classify allergens vs non-allergens, encoding sequence properties with auto- and cross-covariance (ACC), comparing fingerprints by Tanimoto coefficient, and achieving 88% accuracy with a Matthews correlation coefficient (MCC) of 0.759 [51]. Lastly, the ToxinPred2 server was employed to predict the toxicity of the chimeric vaccine.

### Secondary structure and intrinsic disorder prediction

The [PSIPRED v4.0](https://bioinf.cs.ucl.ac.uk/psipred/) and [RaptorX](http://raptorx.uchicago.edu/StructurePropertyPred/predict/) Property web servers were used to accurately predict the secondary structure of the chimeric vaccine based on primary amino acid sequences. The Position-Specific Iterated BLAST (PSI-BLAST) was employed to identify sequences showing significant homology with the vaccine candidates. The performance of PSIPRED was evaluated using a highly stringent cross-validation approach, yielding a Q3 accuracy of 81.6% [52]. RaptorX Property uses the DeepCNF deep-learning framework to predict protein secondary structure, solvent accessibility, and intrinsically disordered regions with high accuracy [53]. Also, the [AIUPred](https://aiupred.elte.hu/) server was employed to assess intrinsically disordered regions of the chimeric vaccine candidate. The AIUPred web server is based on a biophysics-based model enhanced with deep learning techniques [54].

### Prediction of tertiary structure, refinement, and validation

The 3D structure of the designed chimeric vaccine was predicted using [ColabFold](https://colab.research.google.com/github/sokrypton/ColabFold/blob/main/AlphaFold2.ipynb), a user-friendly interface that uses the AlphaFold2 [55] technology within the Google Colab environment. ColabFold offers an enhanced prediction of monomeric protein structures and complexes by combining the fast homology search of MMseqs2 with AlphaFold2 or RoseTTAFold [56]. The obtained 3D structure was further refined using the [GalaxyRefine](https://galaxy.seoklab.org/cgi-bin/submit.cgi?type=REFINE) server. The GalaxyRefine tool improves protein structure quality by rebuilding and repacking side chains, followed by molecular dynamics simulation, achieving superior local quality as demonstrated in CASP10 [57]. The overall quality of the refined 3D structure of the chimeric vaccine candidate was assessed using the [ProSA-web](https://prosa.services.came.sbg.ac.at/prosa.php) server. The ProSA-web tool is commonly used to assess 3D models of protein structures for potential errors and is used for error recognition in experimentally determined structures, theoretical models, and protein engineering [58]. Lastly, the [ERRAT](https://saves.mbi.ucla.edu/) server was also deployed to validate the 3D structure of the chimeric vaccine candidate. The ERRAT server analyzes non-bonded atom-atom interactions compared to reliable high-resolution crystallography structures [59].

### Prediction of discontinuous B-cell epitope

Prediction of conformational B-cell epitopes provides a more accurate representation of antibody-binding sites than linear epitope prediction, offering valuable insights into antigen-antibody interactions [60]. The [ElliPro](https://saves.mbi.ucla.edu/) server was utilized to assess discontinuous B-cell epitopes in the chimeric vaccine. ElliPro identifies and visualizes antibody epitopes based on a modified Thornton approach combined with clustering, MODELLER, and Jmol. In evaluations using conformational epitopes, the tool demonstrated superior performance compared with other predictors, achieving an AUC of 0.732 [61].

### Binding pocket prediction molecular docking, binding affinity, and interaction analyses

To predict the binding pockets in the chimeric vaccine, the [CASTp 3.0](http://sts.bioe.uic.edu/castp/calculation.html) server was used. CASTp is an online structural analysis server that applies an alpha-shape algorithm from computational geometry to identify protein surface pockets and assess their geometric and topological characteristics [62]. Also, to assess the interaction between the designed chimeric vaccine and the innate immune system, molecular docking was performed with Toll-like receptor 4 (TLR4) using the [HawkDock](https://cadd.zju.edu.cn/hawkdock/) server. The refined 3D structures of the vaccine candidate and TLR4 were submitted to HawkDock as ligand and receptor, respectively, for the docking calculations. The HawkDock server, an integrated platform for protein-protein docking, was used to evaluate interactions between the vaccine construct and TLR4. The server combines molecular docking with molecular mechanics/generalized Born surface area (MM/GBSA) calculations to estimate binding free energies. Among the generated docking models, the complex with the lowest MM/GBSA binding free energy was selected, as it is considered the most thermodynamically stable and representative of a native-like protein-protein interaction [63, 64]. The selected docked complex was subsequently visualized using PyMOL.

The binding affinity of the interaction between the vaccine candidate and the TLR4 immune receptor was analyzed using the PROtein binDIng enerGY ([PRODIGY](https://wenmr.science.uu.nl/login)) prediction server. The server implements a straightforward but very efficient predictive model based on intermolecular contacts and properties derived from the non-interface surface [65]. The docked complexes were then further analyzed on the [PDBsum](https://www.ebi.ac.uk/thornton-srv/databases/pdbsum/) server, which creates a 2D image of the interactions at the ligand-protein interface from the 3D coordinates.

### Normal mode analysis

Normal mode analysis (NMA) for the docked chimeric vaccine candidate-TLR4 complex was assessed using the [iMODS](https://imods.iqf.csic.es/) server. The analysis was performed to examine the internal dihedral coordinates and evaluate the cooperative functional motions of the docked complex. The essential dynamic simulation program of the iMODS server was used to ensure energy minimization, molecular stability, and atomic mobility of the chimeric vaccine in the docked complex. The server was used to analyze the possible motions in the vaccine candidate-TLR4 complex based on specialized parameters, including B-factors, RMSD, eigenvalues, deformability of the complex, covariance values, and elastic models [66].

### Codon optimization and *in silico* cloning

Reverse translation of the vaccine protein sequence into a DNA sequence and codon optimization were done using the [Java Codon Adaptation Tool](https://www.jcat.de/) (JCat) server. JCat optimizes codon usage to match the preferred codon bias of a selected host organism, thereby enhancing the potential for efficient heterologous protein expression. In addition, the server calculates the codon adaptation index (CAI) and GC content, both of which are important determinants of gene expression efficiency in bacterial hosts [67, 68]. To express the designed chimera in *E. coli,* the codon-optimized gene sequence was inserted between the *Nco*I and *SgrA* I restriction sites of the pET30a+ vector using the [SnapGene](https://www.snapgene.com/) tool.

### Immune simulation

To further predict the immunogenicity and immune response profile of the chimeric vaccine, *in silico* immune simulations were conducted using the C-ImmSim server as previously described [22]. C-ImmSim is an agent-based immune simulation platform that uses position-specific scoring matrices (PSSM) and machine-learning approaches to predict epitopes and model immune interactions. It simulates key mammalian immune compartments, including the bone marrow, thymus, and a tertiary lymphoid organ, to represent hematopoiesis, T-cell selection, and immune responses, respectively [69]. All simulation parameters were maintained at their default settings. Vaccine injections were simulated at time steps 0, 42, and 84, corresponding to two weeks apart. Each time step corresponded to 8 hours, with time step 0 representing the initial injection, as previously described by [22].

## Results

### Protein selection for vaccine design

The amino acid sequences of the 03 proteins (*Na*-APR-1, *Na*-GST-1, and *Na*-ASP-2), previously reported to be immunogenic, implicated in pathogenesis or virulence in *N. americanus* (accession numbers: Q9N9H3, D3U1A5 and Q7Z1H1) were retrieved from the UniProtKB database for preliminary analyses, epitope prediction and design of a chimeric antigen, while the tertiary structure of human TLR4 agonist (PDB ID: 4G8A) was obtained from the [Protein Data Bank](https://www.rcsb.org/). The TLR4 agonist used in this study is a 50S ribosomal protein from *M. tuberculosis*, known to activate TLR4 signaling, and was used as a molecular control adjuvant in the docking and simulation analyses [70].

### Signal peptide, sequence localization, and transmembrane domain prediction

Both SignalP-6.0 and TOPCONS servers predicted the presence of signal peptides only in *Na*-APR-1, while no signal peptides were predicted for either *Na*-GST-1 or *Na*ASP-2. For TM domain prediction, all three proteins were predicted by both DeepTMHMM 1.0 and TOPCONS to contain no transmembrane domains. Also, both the DeepLoc 2.1 and WoLF PSORT servers predicted *Na*-ASP-2 to be localized in the extracellular space. On the other hand, *Na*-GST-1was predicted to be localized in the cytoplasm by both servers, while *Na*-APR-1 was predicted to be localized in the Lysosome/Vacuole by the DeepLoc 2.1 server and the plasma membrane by the WoLF PSORT server (**Table 1**).

**Table 1:** Predicted signal peptides, TM domain and protein localization.

| Protein | Signal peptide |  | TM domain |  | Localization |  |
| --- | --- | --- | --- | --- | --- | --- |
|  | SignalP 6.0 | TOPCONS | TOPCONS | DeepTMHMM 1.0 | DeepLoc 2.1 | WoLF PSORT |
| *Na*-APR-1 | Yes | Yes | No | No | Lysosome/Vacuole | Plasma membrane |
| *Na*-GST-1 | No | No | No | No | Cytoplasm | Cytoplasm, |
| *Na*-ASP-2 | No | Yes | No | No | Extracellular | Extracellular |

### Analysis of protein conservation across hookworm species

A significant degree of conservation among all selected proteins was noted in related species of the hookworm species, which ranged from 48.8 % to 100%, based on a BLAST search conducted against the UniProtKB database. The analysis of protein conservation compared to their homologs in *A. caninum, A. ceylanicum,* A*. duodenale* and *Haemonchus contortus* revealed that *Ancylostoma duodenale* showed the highest level of conservation, with *Na*-APR-1 protein showing 91.3% conservation, while *Haemonchus contortus* had the lowest level of conservation, with *Na*-ASP-2 showing 58.6% (**Table 2**).

**Table 2:** Conservation of target proteins in selected *hookworms*.

| Number | Protein | Percentage identity (%) |  |  |  |
| --- | --- | --- | --- | --- | --- |
|  |  | *Ancylostoma duodenale* | *Ancylostoma caninum* (Dog hookworm) | *Ancylostoma ceylanicum* | *Haemonchus contortus* |
| 1 | *Na*-APR-1 | 91.3 | 90.1 | 89.4 | 79.5 |
| 2 | *Na*-GST-1 | 78.2 | 69.6 | 81.6 | 66.2 |
| 3 | *Na*-ASP-2 | 63.6 | 61 | 58.6 | 58.6 |

### Prediction of cytotoxic T lymphocytes (CTL) epitopes

High-scoring CD8+ (9-mer) epitopes were identified from the antigens using the NetCTL 1.2 server, at the default threshold of 0.75. From the predicted epitope sequences, eight CD8+ epitopes were selected for incorporation in the chimera based on their antigenicity (**Table 3**).

**Table 3:**
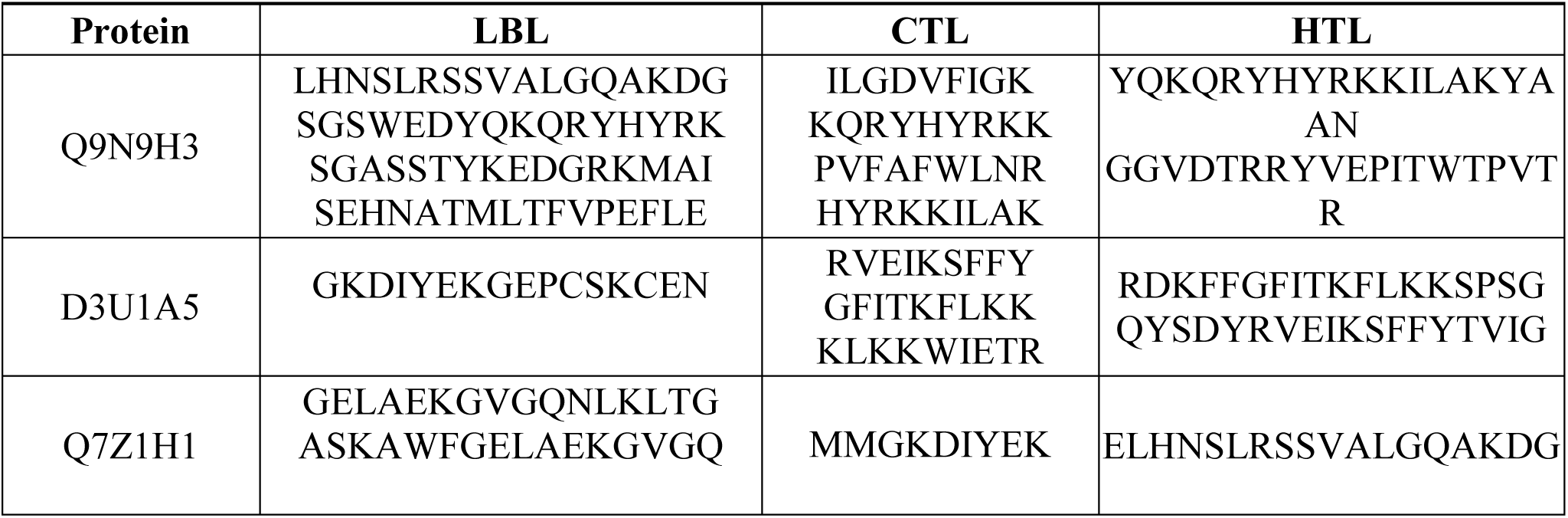
Epitopes predicted for the design of the multi-epitope vaccine.

### Prediction of Helper T-cell (HTL) epitopes

High-affinity MHC-II epitopes, predicted based on IC50 scores for the HLA-DR, HLA-DQ, and HLA-DP human alleles, were predicted using the NetMHCII 2.3 web server and identified as HTL epitopes. Six high-affinity HTL epitopes were selected for the design of the chimeric vaccine candidate. Some of the predicted CTL epitopes and HTL epitopes were found to have overlapping regions (**Table 3**).

### Prediction of linear B-lymphocyte (LBL) epitopes

16-mers of linear B-cell epitopes were predicted using ABCpred, and those identified as highly antigenic were chosen for constructing the chimeric vaccine candidate. In total, seven linear B-cell epitopes were chosen to be incorporated into the chimera (**Table 3**).

### Design of the chimeric vaccine candidate

The predicted epitopes were combined using different linker types (AAY, GPGPG, and KK linkers to connect CD8+ T-epitopes, CD4+ T-epitopes, and linear B-cell epitopes, respectively) to form a chimeric vaccine (designated EpiNecaVac). The designed chimeric vaccine candidate consisted of 08 CTL epitopes, 05 HTL epitopes, and 07 linear B-lymphocyte epitopes in total. In addition to the predicted epitopes, the TLR4 agonist from the *M. tuberculosis* 50S ribosomal protein L7/L12 (Locus RL7_MYCTU, UniProt ID: P9WHE3) as chosen as a built-in adjuvant and incorporated at the N-terminus of the designed vaccine candidate using the EAAAK linker to enhance the elicited immune responses. The 6xHis tag was added to the carboxyl terminus of the designed chimeric antigen to facilitate purification and identification. In summary, the EpiNecaVac chimeric vaccine candidate consisted of 481 amino acid residues, which include 20 peptides from *N. americanus* (predicted epitopes), as well as an immune enhancer (TLR4 agonist) and various linkers (**Figure 2**).

**Figure 2.**
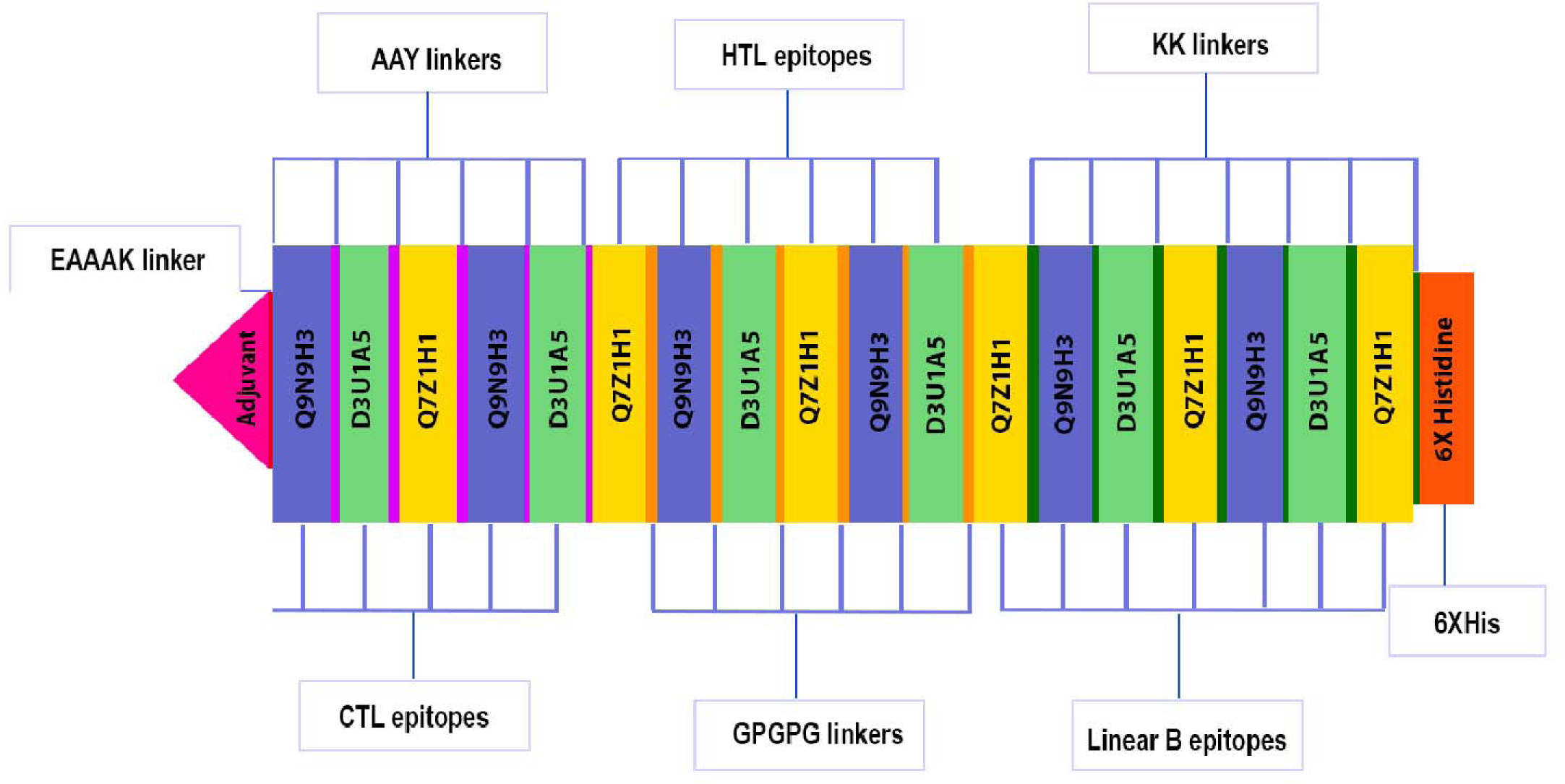
Schematic illustration of the final multi-epitope vaccine antigen EpiNecaVac. This 481-amino acid polypeptide sequence includes a built-in adjuvant (pink) at the N-terminus, linked to the multi-epitope sequence via an EAAAK linker (red),the antigenic CTL, HTL, and linear B cell epitopes are joined by AAY (purple), GPGPG (orange), and KK linkers (green), respectively. A 6x-His tag (orange), is incorporated at the C-terminus for purification and identification, positioned away from the peptide.

### Analysis of physicochemical properties and solubility

The EpiNecaVac chimeric antigen, composed of 481 amino acid residues, was predicted to have a molecular weight (MW) of 52.9 kDa and a theoretical pI of 9.68, indicating that the protein is basic. The protein was predicted to have a half-life of 30 hours in mammalian reticulocytes *in vitro*, > 20 hours in yeast, and >10 hours in *E. coli in vivo*. Furthermore, physicochemical property analyses showed the protein to be rich in Lys (14.1%) and Ala (13.1%). The predicted instability index (II) was calculated to be 28.88. A protein with an instability index <40 is expected to be stable, while a value >40 suggests that the protein may be unstable [46]. The predicted GRand Average of hydropathicitY (GRAVY) was -0.541, suggesting that the protein is hydrophilic and capable of interacting with water molecules [71]. An aliphatic index of 69.92 was predicted, implying that the protein exhibits thermostability [72]. The solubility score was predicted to be 0.8158 by the NetSolP 1.0 server and 0.5757 on the DeepSoluE server. Both servers predicted the protein to be soluble upon expression. Furthermore, the NetSolP 1.0 server predicted the protein to have a usability score of 0.4229 (**Table 5**).

**Table 5:**
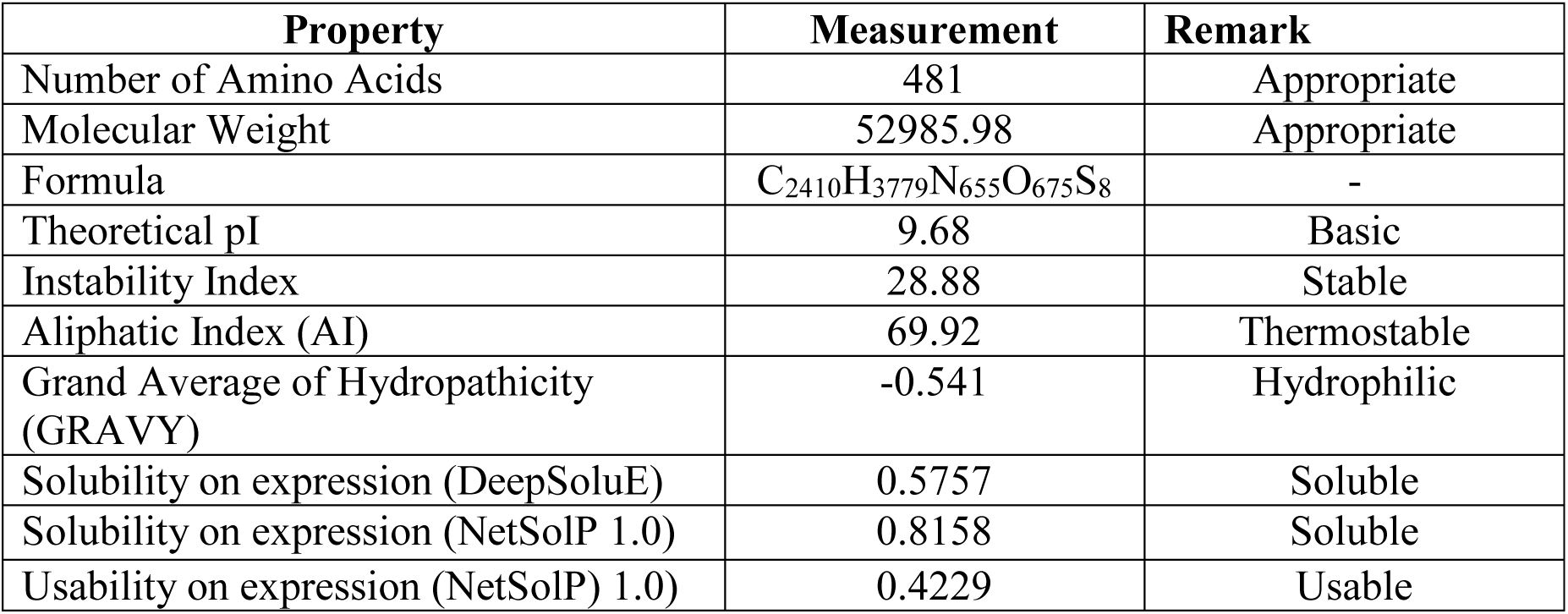
Physicochemical properties of the designed chimeric vaccine.

### Secondary structure and intrinsic disorder prediction

The secondary structure of the chimeric vaccine was predicted to consist of 44% alpha helices, 7% beta-strands, and 49% coils (**Figure 3A**). Concerning solvent accessibility, EpiNecaVac was predicted to have 58% of amino acid residues exposed, 20% fairly exposed, and about 21% were predicted to be buried. A total of 178 amino acid residues, accounting for 37%, were predicted to be found in intrinsically disordered regions, while 63% were predicted to be ordered (**Figure 3B**). Figure **3C** presents a visual representation of the secondary structure of the designed vaccine candidate.

**Figure 3.**
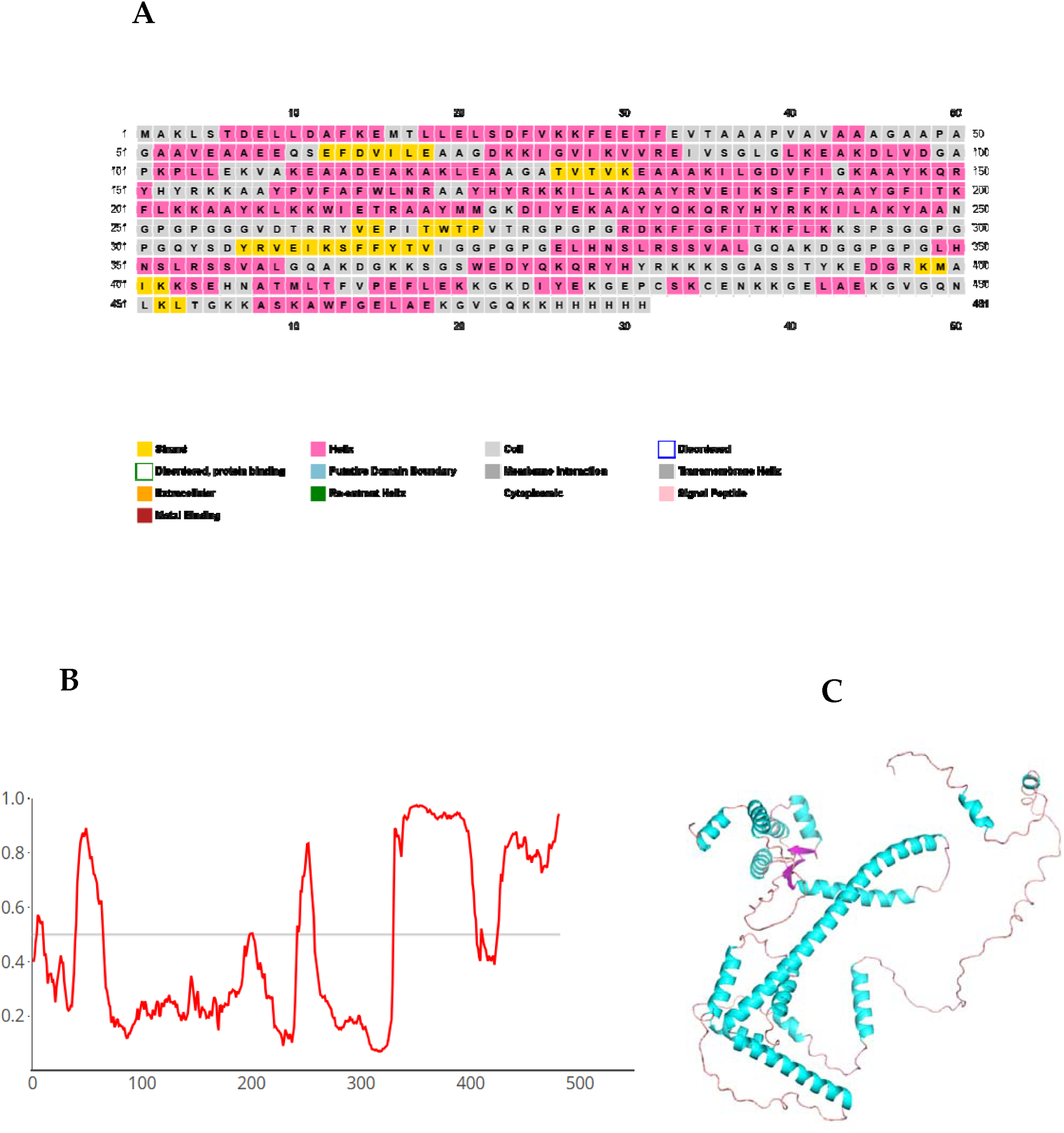
Graphical representation of the secondary structure of the final subunit vaccine sequence. (A) The protein is predicted to consist of alpha-helices (44%), beta strands (7%), and coils (49%). (B) According to AIUpred, 37% of positions are expected to be in intrinsically disordered regions. (C) The 3D structure of the vaccine candidate displays the various secondary structure elements, with helices in cyan, strands in pink, and coils in orange.

### Prediction of antigenicity, allergenicity, and toxigenicity of the chimeric vaccine

The final sequence had a predicted antigenicity score of 0.7987 on the VaxiJen 2.0 server, using the parasite model with a threshold of 0.5. An antigenic score of 0.7716 was predicted for the designed chimeric construct on the ANTIGENpro server, suggesting it is likely a vaccine candidate. The AllerTOP v.2 and AllergenFP server’s prediction showed that the protein is non-allergenic. A non-allergenic prediction suggests that the vaccine candidate is unlikely to trigger an allergic response in most individuals, making it safer for widespread use [73]. The ToxinPred2 server predicted that EpiNecaVac did not have any toxigenic peptides (**Table 6**).

**Table 6:** Antigenicity, allergenicity, and toxigenicity prediction of the designed vaccine.

| Property | Prediction | Remark |
| --- | --- | --- |
| Antigenicity (VaxiJen v2.0) | 0.7987 | Probable antigen |
| Antigenicity (ANTIGENpro) | 0.7716 | Probable antigen |
| Allergenicity (AllerTOP v2.0) | Probable NON-ALLERGEN | Probable NON-ALLERGEN |
| Allergenicity (AllergenFP 2.0) | Probable NON-ALLERGEN | Probable NON-ALLERGEN |
| Toxicity (ToxinPred2) | Non-toxin | Non-toxigenic |

### Prediction of epitopes for mouse MHC II alleles

Out of the 481 potential 9-mer epitopes derived from the antigen for each allele, 156 peptides were predicted to bind to six mouse MHC II alleles for EpiNecaVac. Among these, the H-2-IEd allele showed the highest number of epitopes, while the H-2-IAk and H-2-IEk alleles had the lowest binding epitopes (**Table 7**).

**Table 7:** Predicted peptide counts for binding to mouse H2 class II alleles.

| Allele | H-2-IAb | H-2-IAd | H-2-IAk | H-2-IAs | H-2-IAu | H-2-IEd | H-2-IEk |
| --- | --- | --- | --- | --- | --- | --- | --- |
| Number of epitopes | 25 | 17 | 2 | 19 | 35 | 56 | 2 |

### Prediction of IFN-γ-inducing epitopes

Out of the 473 potential IFN-γ-inducing epitopes (15-mers) predicted for the chimeric vaccine, 176 exceeded the epitope prediction threshold. The predicted quantity of these IFN-γ-inducing epitopes was correlated with the simulated levels of IFN-γ induction observed after immunization with the chimeric vaccine, as modeled by immune simulation on the C-ImmSim server (**Figure 4**).

**Figure 4.**
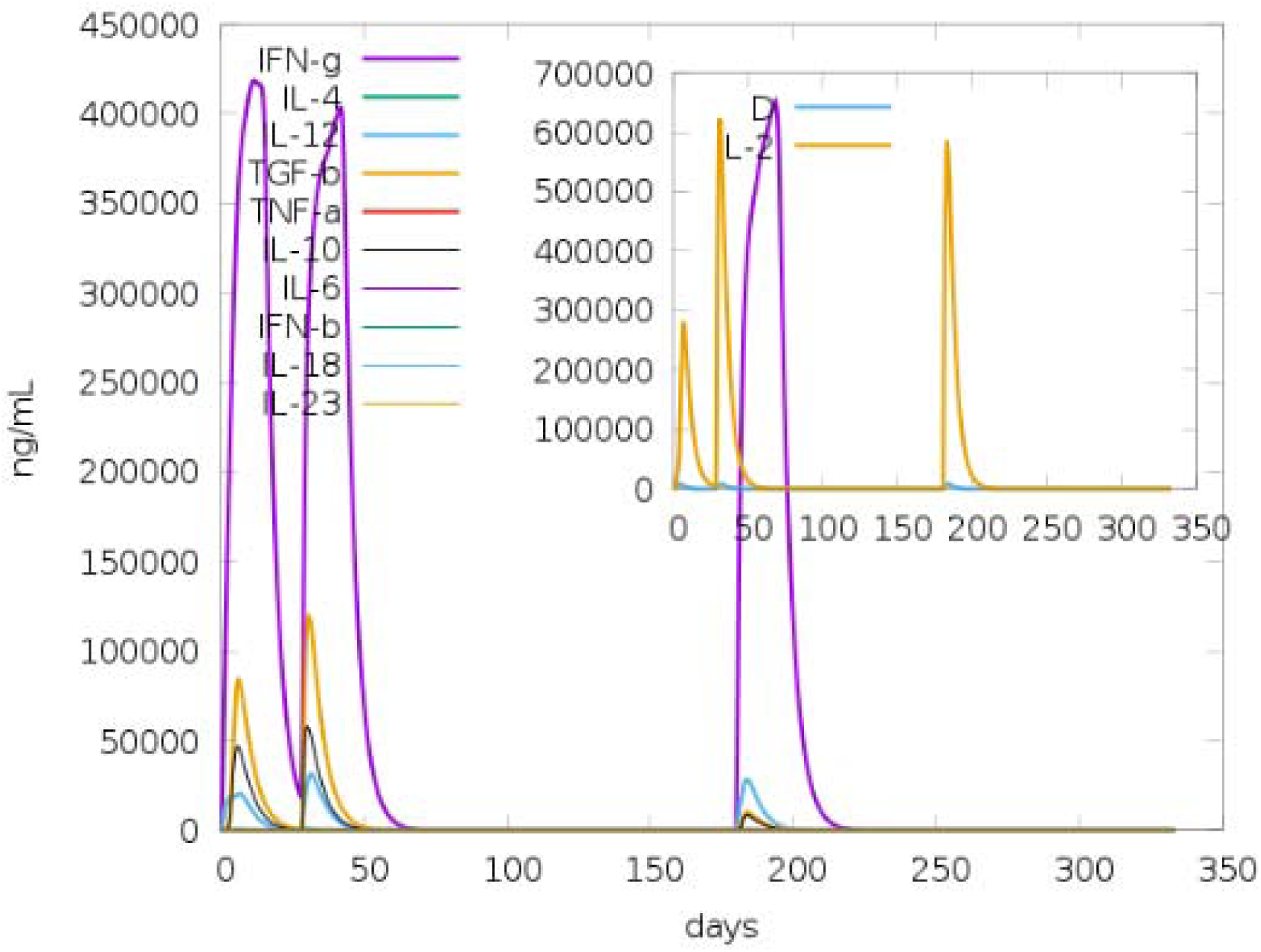
Immune simulation of the cytokine profile induced by three injections of EpiNecaVac administered three days apart is presented. The main plot illustrates the varying cytokine levels following the three injections, while the inset plot displays the IL-2 level along with the Simpson index, D, represented by the blue line. The index D measures diversity, with an increase over time indicating the emergence of different epitope-specific dominant T-cell clones. A smaller D value corresponds to lower diversity.

### Prediction of tertiary structure, refinement, and validation

The 3D structure of the multi-epitope vaccine candidate was predicted using AlphaFold2 on the ColabFold interface (**Figure 5A**). 3D structural refinement was done on the GalaxyRefine server, which yielded 5 models. Among these 5 models, “model 4” was selected with parameters GDT-HA (0.8867), RMSD (0.650), MolProbity (0.901), and Rama favored (97.9), which was then used for downstream analyses (**Figure 5B**). In addition, the model was evaluated to have a clash score of 1.4, a poor rotamers score of 0, and a Ramachandran plot score of 95.9%. The analysis of the Ramachandran plot of the selected model showed 95.9% of protein residues to be located in favored regions, which is similar to what was obtained from the GalaxyRefine server. Furthermore, it was predicted that 3.5% of residues were predicted to be located in allowed regions, while only 0.2% were predicted in disallowed regions (**Figure 5C**). Following refinement, the quality of the 3D model was analyzed on the ERRAT and ProSA-web servers. The selected model has an overall quality factor of 90.2 on the ERRAT server and a Z-score of -3.26 on the ProSA-web server (**Figure 5D**). A ProSA-web Z-score of -3.26 suggests the protein model is likely within the typical range for native proteins of similar size, indicating reasonably good quality, but further investigation is recommended to identify potential localized issues [58].

**FIGURE 5.**
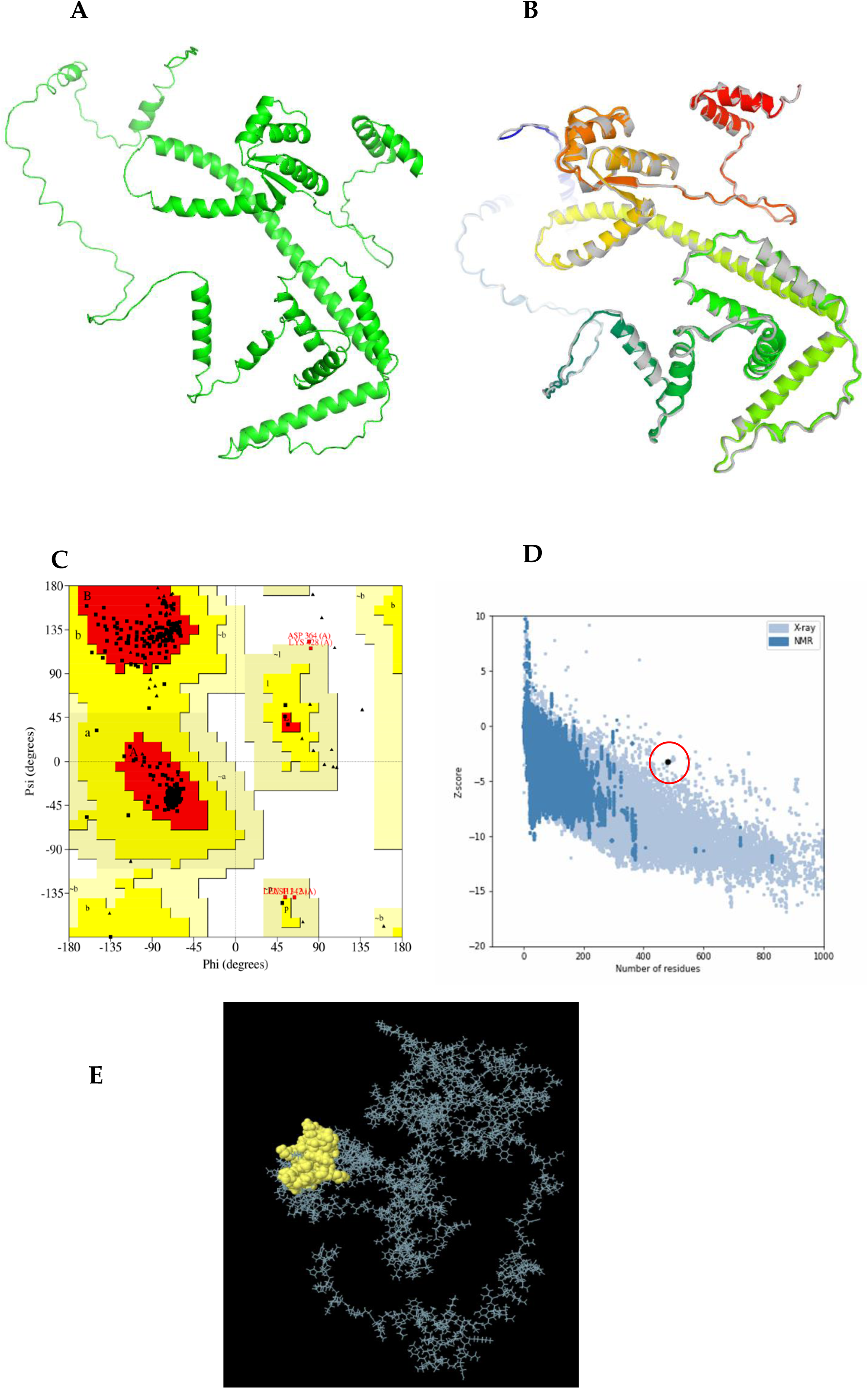
Protein modeling, refinement, and validation: (A) the initial 3D model of the multi-epitope vaccine candidate was predicted using the AlphaFold2 server. (B) The refined 3D structure (colored) was superimposed on the initial "crude model" (gray) using the GalaxyRefine server. Validation of the refined model revealed (C) A Ramachandran plot analysis showing that 95.9% of protein residues (green crosses) are in favored regions, 3.2% (orange triangles) in allowed regions, and 0.7% (red circles) in disallowed (outlier) regions. Additionally, (D) ProSA-web provided a Z-score of -3.26, and (E) discontinuous B-cell epitopes are indicated in yellow on the chimeric vaccine construct.

### Prediction of discontinuous B-lymphocyte (LBL) epitopes

In total, there were 24 residues (42%) predicted by the ElliPro server to be found in 24 conformational B-cell epitopes, with predicted epitope scores in the range of 0.576 to 0.976. The predicted discontinuous epitope residues ranged from 3 to 28 amino acid residues (**S1 Table 1**.) and (**Figure 5E**).

### Protein-protein interaction between the TLR4 Receptor and the chimeric vaccine

TLR4 has been implicated in protective immune responses against helminth infections, including *O. volvulus* and *S. stercoralis* [74, 75], while TLR4 agonists have also been investigated as adjuvants in *N. americanus* vaccine development [76]. Molecular docking investigated the binding interaction between the refined and validated 3D structure of the chimeric vaccine candidate and TLR4 after a protein-protein interaction pocket was predicted in the designed chimera (**Figure 6 A**). One hundred models were generated and visualized, with model 1 selected for downstream analysis. The selected EpiNecaVac-TLR4 complex model exhibited the lowest energy of -5192.69 (kcal/mol) with an MM/GBSA value of -63.27 (kcal/mol) (**Figure 6 B**).

**FIGURE 6.**
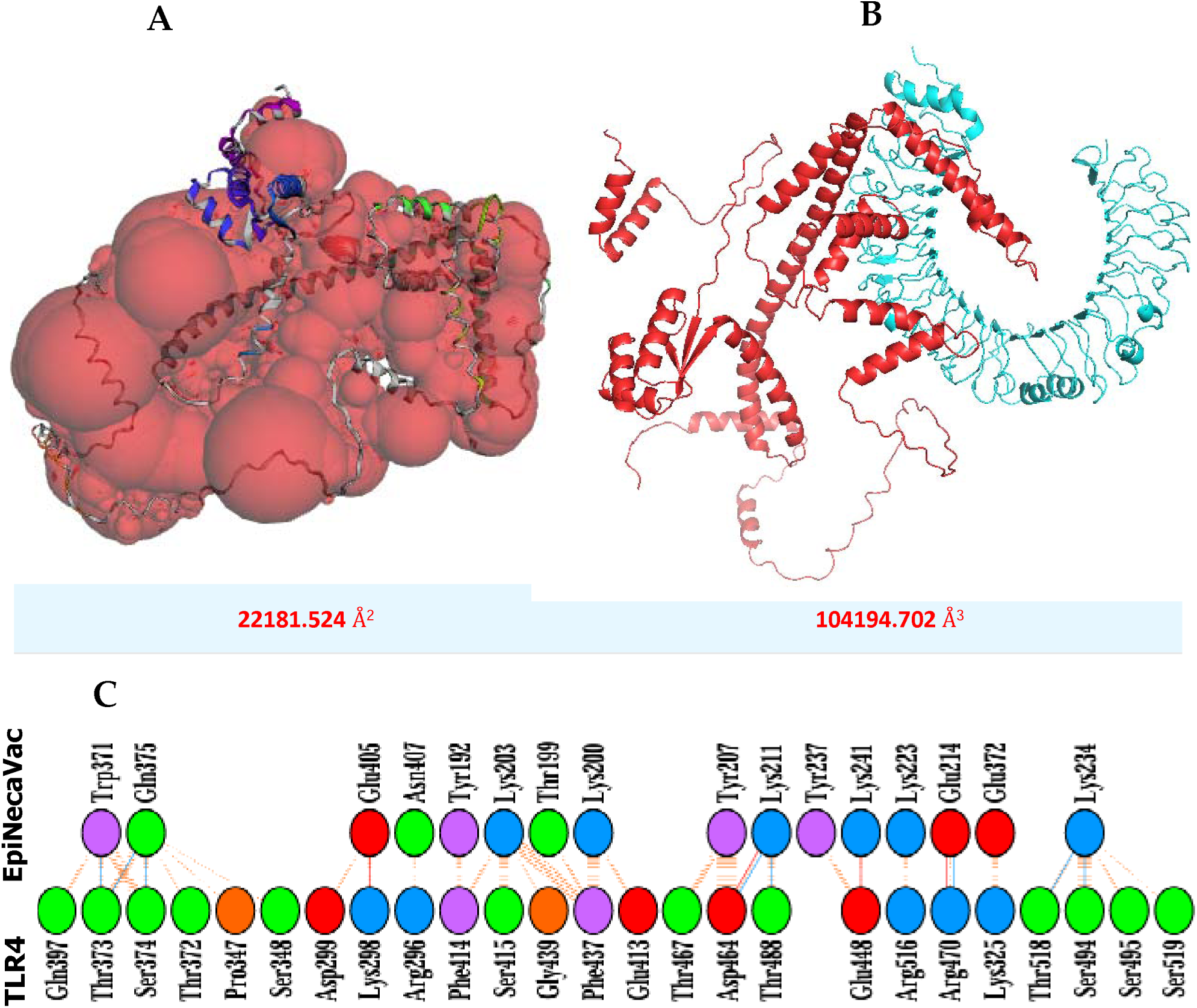
Protein-protein interaction pocket prediction and molecular docking of the vaccine candidate with the TLR4 receptor were conducted. (**A**) The largest predicted PPI pocket is located at the interface of the designed chimeric vaccine candidate (in red), with a surface area of 22181.524 Å² and a surface volume of 104194.702 Å³. (**B**)The docked complex of the designed vaccine candidate (red) and the Toll-like receptor 4 chain (cyan) is shown. (**C**) The interaction network between the vaccine candidate and TLR4 is illustrated, with hydrogen bonding interactions depicted in blue. Non-bonded interactions and salt bridges are represented in orange. Charged residues are color-coded: positive in blue, negative in red, neutral in green, aliphatic in grey, and aromatic in purple.

The PDBsum server predicted the formation of 8 hydrogen bonds, 4 salt bridges, and 151 non-bonded contacts between 25 residues from TLR4 and 16 residues of the chimeric vaccine candidate (**Figure 6 C**). Overall, the vaccine candidate demonstrated favorable interactions with TLR4. The relative binding free energy (ΔG) of the EpiNecaVac-TLR4 complex was predicted to be -10. Kcal/mol, while the predicted Kd was 3.1e-08. Consequently, the number of contacts made at the interface (IC) per property (ICs charged-charged: 12.0, ICs charged-polar: 23.0, ICs charged-apolar: 13.0, ICs polar-polar: 4.0, polar-apolar: 17.0, ICs apolar-apolar: 3.0).

### Normal mode analysis

The deformability graph revealed peak points that signify the deformed regions of the main chain residues in the vaccine candidate-TLR4 complex. The areas identified as hinges correspond to regions of high deformability (**Figure 7A**). The B-factor plot illustrates the relationship between NMA mobility and the EpiNecaVac-TLR4 complex, showing the average RMSD values of the docked complex (**Figure 7B**). B-factor values indicate the uncertainty associated with each atom. The computed eigenvalue for the multi-epitope vaccine candidate-TLR4 complex was 7.974841e-08, which reflects motion stiffness with respect to each normal mode (**Figure 7C**). The eigenvalue estimates the energy needed to deform the structure. Each normal mode of the complex is depicted by individual (purple) and cumulative (green) variance in the variance bar. It is worth noting that there is a negative correlation between the variance and the eigenvalue (**Figure 7D**). Furthermore, the interactions between the multi-epitope vaccine candidate and TLR4 in the complex are depicted using a covariance matrix. The related motions among various pairs of residues are indicated by correlated (red), uncorrelated (white), and anti-correlated (blue) atomic movements within the multi-epitope vaccine candidate-TLR4 complex (**Figure 7E**). An elastic network map was also created to show atom pairs linked by springs in the multi-epitope vaccine candidate-TLR4 complex. Each dot in the graph denotes a spring connecting the corresponding pair of atoms. The color of the dots indicates their stiffness; darker shades of grey represent stiffer regions, while lighter dots signify more flexible areas (**Figure 7F**). All the findings from the normal mode analyses suggest the favourable interaction and stability within the multi-epitope vaccine candidate-TLR4 complex.

**Figure 7.**
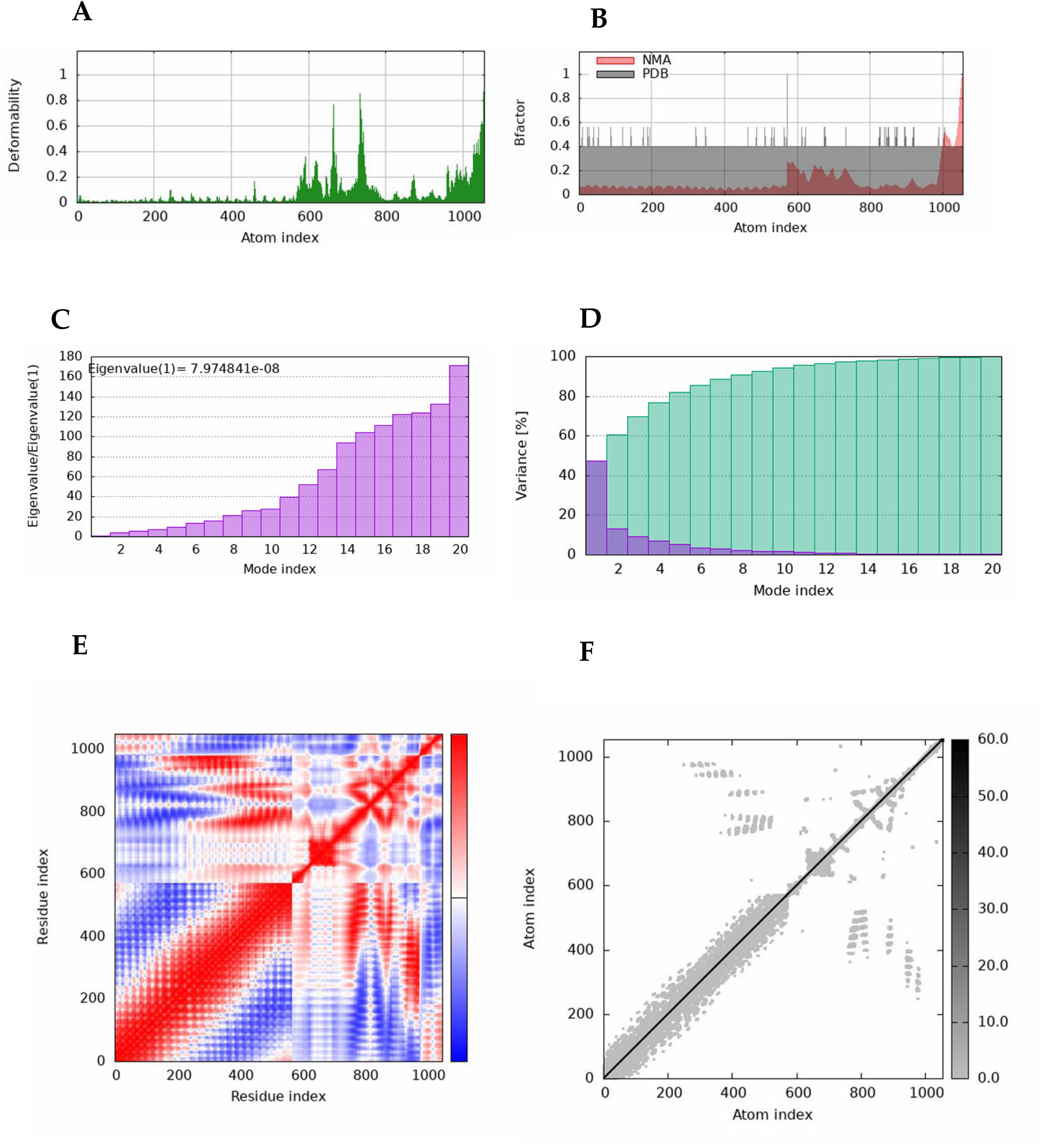
Normal mode analysis of the vaccine candidate-TLR4 docked complex. (A) Displays the main-chain deformability of the EpiNecaVac-TLR4 complex. (B) The B-factor quantifies the uncertainty associated with each atom. (C) The eigenvalue indicates the motion stiffness related to each normal mode. (D) The variance map shows individual (red) and cumulative (green) variances. (E) The covariance graph illustrates the mobility of residue pairs, indicating correlated (red), uncorrelated (white), or anti-correlated (blue) movements. (F) The elastic network model represents pairs of atoms connected by springs, with darker greys indicating greater stiffness of the springs.

### Immune simulation analysis

The primary immune response on the C-ImmSim server was characterized by high levels of IgM. The secondary and tertiary responses exhibited significant increases in B-cell populations and in the levels of IgG1, IgG2, IgM, and the combined IgG and IgM antibodies, accompanied by a corresponding decrease in antigen concentration (**Figure 8A, B**). This profile suggests the development of immune memory, leading to enhanced clearance of the antigen during subsequent exposures (**Figure 8C**). A similarly strong response was observed in the Th (helper) and TC (cytotoxic) cell populations, accompanied by corresponding memory development (**Figure 8C, D**). Repeated exposure with 03 injections (administered two weeks apart) triggered increasing levels of IgG1 and decreasing levels of IgM, while IFN-γ concentration and the TH cell population were maintained at high levels throughout the duration of exposure. In summary, the developed vaccine candidate may induced both cellular and humoral immune responses, which are recognized as crucial for protection against *N. americanus*.

**Figure 8.**
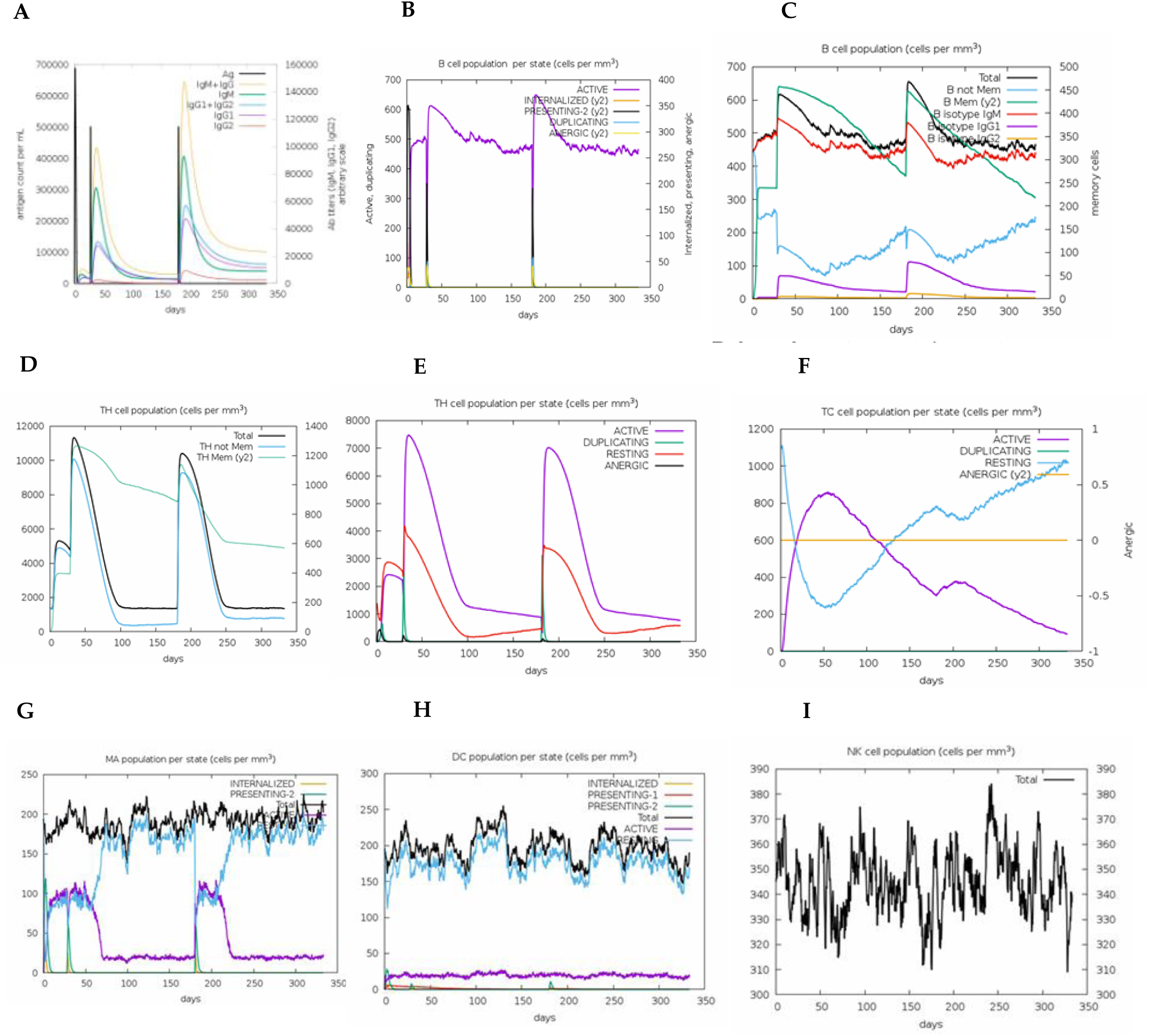
In silico simulation of the immune response triggered by the designed vaccine candidate: (**A**) Antigen and immunoglobulins, (**B**) B cell population by state, (**C**) Total B cell population, (**D**) TH cell population, (**E**) TH cell population by state, (**F**) TC cell population by state, (**G**) MA population by state, (**H**) DC population by state, and (**I**) NK cell population.

### Population coverage analysis

The overall coverage for MHC class I CTL epitopes was 63.96%, while MHC class II HTL epitopes showed a coverage of 100% (**Fig. 9A and B**). For specific regions, the predicted coverage for MHC class I CTL epitopes ranges from 41.61% to 66.11%, with South America having the highest percentage coverage while South Africa has the lowest. Interestingly, for MHC class II HTL epitopes, all regions had 100% coverage, except for South Africa, which also had the lowest predicted percentage coverage (72.44%) (**S1 Fig. 1 and 2**). This detailed regional analysis allows for a better understanding of the vaccine’s potential efficacy in diverse populations across these areas, highlighting the importance of targeting regions with higher predicted coverage. This information is vital for guiding future clinical trial designs and vaccine deployment strategies.

**Figure 9.**
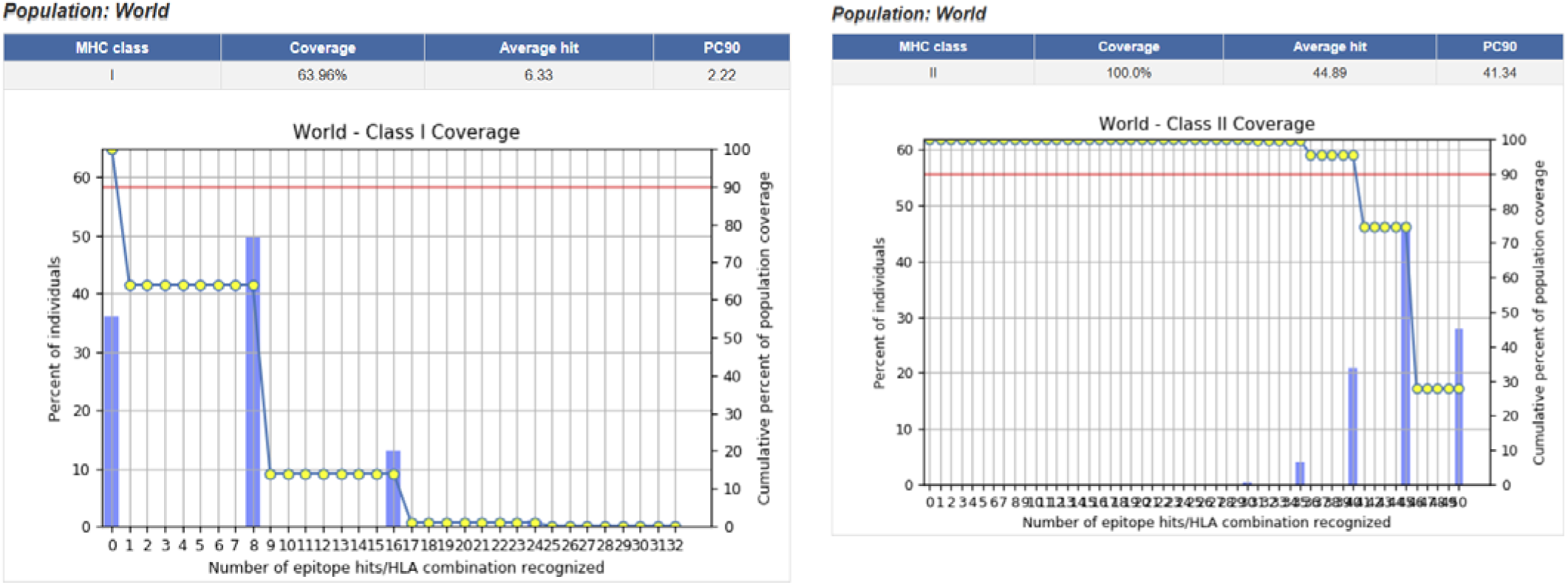
Population coverage analysis of predicted T-cell epitopes across the global population is presented as follows: (A) MHC class I epitope coverage demonstrates a global population coverage of 63.96%, with average hit and PC90 values suggesting extensive HLA representation. (B) MHC class II epitope coverage reflects a global population coverage of 100%, highlighting the distribution of HLA class II alleles across various populations. The histograms and cumulative curves illustrate the percentage of individuals recognizing a specific number of epitopes-HLA combinations.

### In silico cloning

The gene encoding the designed chimeric vaccine candidate was codon-optimized using the Java Codon Adaptation Tool (JCat) to enhance protein expression levels in *E. coli*. The total gene sequence comprised 1444 nucleotides, featuring a Codon Adaptation Index (CAI) of 1.0 and an average GC content of 47.3%. These findings indicate a favorable potential for expression in the *E. coli* bacterial host. Typically, a GC content between 30% and 70% is considered optimal for effective protein expression [77, 78]. Finally, the adapted codon sequence was inserted into the appropriate sites of the pET30a (+) vector using SnapGene software v3.2.1 to create a recombinant plasmid *in silico* (**Figure 10**).

**Figure 10.**
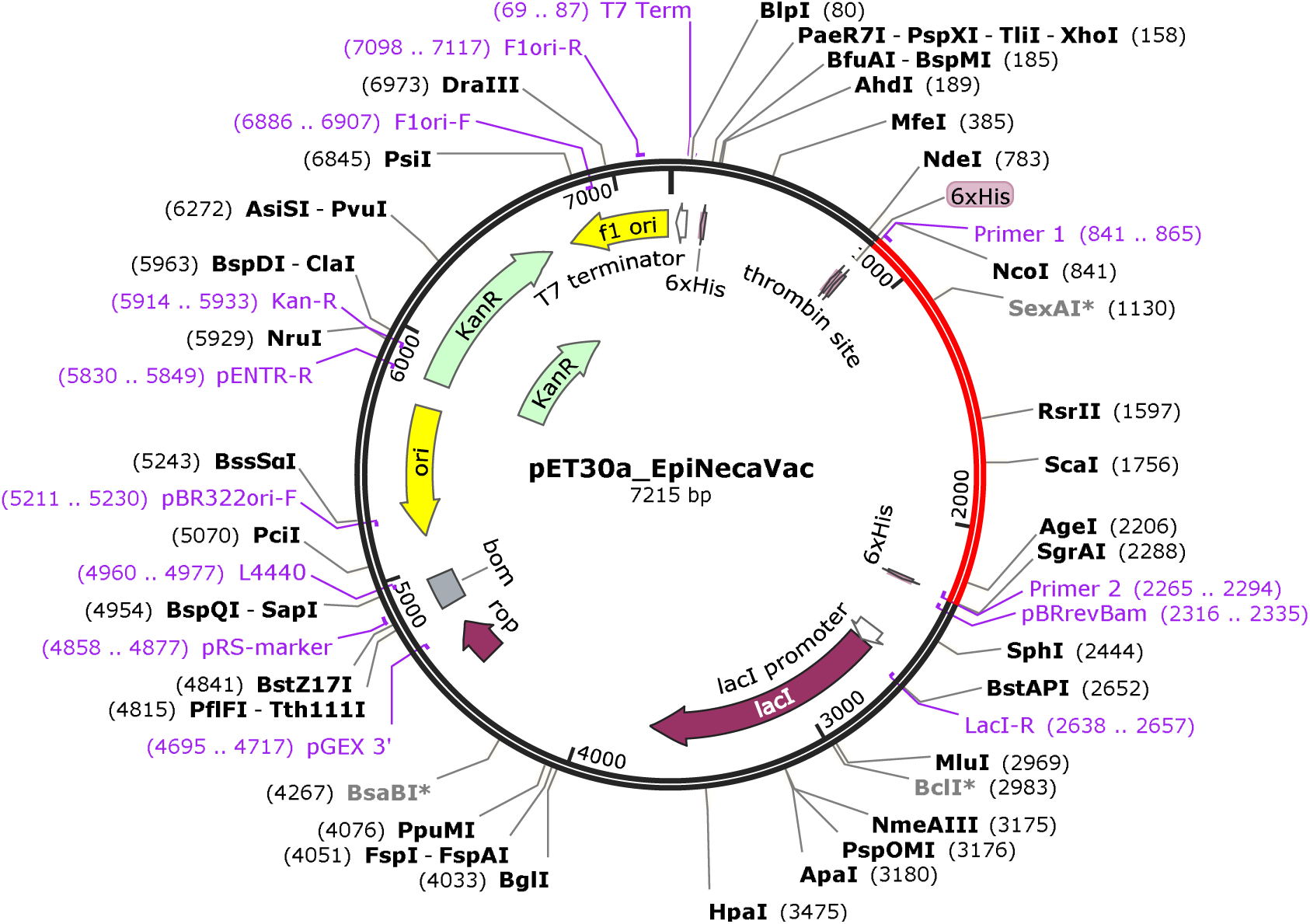
In silico restriction cloning of the final vaccine sequence into the pET30a (+) expression vector is shown, with the red section indicating the gene coding for the vaccine and the black circle representing the vector backbone.

## Discussion

Neglected tropical diseases (NTDs) are a group of infectious diseases that disproportionately affect populations in tropical and subtropical regions, particularly marginalized communities with limited access to healthcare, safe drinking water, and adequate sanitation [7]. Hookworm infection caused by *Necator americanus* remains an important neglected parasitic disease, and although several potential vaccine antigens have been identified, only a limited number have progressed to advanced preclinical and clinical evaluation [8–10]. Although chemotherapeutic interventions are available for many NTDs and hookworm in particular, their long-term effectiveness is limited by frequent reinfection, the emergence of drug resistance, and challenges associated with sustained implementation of treatment programs. Therefore, the development of preventive and/or therapeutic vaccines against NTDs is essential for achieving sustainable and long-term disease control [79, 80]. Indeed, recent modelling studies suggest that preventive chemotherapy alone is unlikely to be sufficient to interrupt hookworm transmission and would be most effective when combined with complementary public-health interventions [81]. Hookworm vaccine development efforts have predominantly focused on three subunit antigens targeting distinct parasite stages, each with promising progress but specific limitations that remain unresolved. Epitope-based vaccines provide a new strategy for eliciting targeted immune responses while preventing reactions to unwanted epitopes in the entire antigen [82]. Epitope-based vaccines offer improved safety, enhanced immunogenicity, and targeted responses against conserved epitopes [83].

The present study designed and computationally validated EpiNecaVac, a chimeric vaccine candidate against *N. americanus*, constructed from carefully filtered B-cell, CTL, and HTL epitopes drawn from the three clinical-stage hookworm antigens *Na*-APR-1, *Na*-GST-1, and *Na*-ASP-2, together with an *M. tuberculosis* 50S ribosomal protein L7/L12 TLR4-agonist adjuvant. This design directly addresses one of the central unresolved problems in hookworm vaccinology identified in the introduction: no single recombinant antigen combines B-cell and T-cell epitope breadth with a safety profile suitable for populations already exposed to natural infection. By computationally excising the IgE-reactive and toxic determinants of *Na*-ASP-2 while retaining its antigenic epitopes, the present approach aims to recover the immunogenic value of an antigen whose clinical development was previously halted due to allergenicity, without reintroducing the associated risk.

*Na*-ASP-2 was predicted to be extracellular, and none of the three parent proteins carried predicted transmembrane domains, supporting their suitability as accessible, epitope-rich targets [84]. Cross-species conservation varied by antigen, with *Na*-APR-1 showing the highest identity to its orthologs in *Ancylostoma duodenale,* followed by *Na*-GST-1, while *Na*-ASP-2 was the least conserved (58.6-63.6%). This gradient suggests that epitopes derived from *Na*-APR-1, and to a lesser extent *Na*-GST-1, carry the greatest potential for cross-protection against related hookworm and nematode species, though cross-protection remains a prediction requiring confirmation in heterologous challenge studies.

The refined model met conventional structural benchmarks: 95.9% of residues in favoured Ramachandran regions, an ERRAT quality factor of 90.2, and acceptable GalaxyRefine metrics (GDT-HA 0.8867, RMSD 0.650). The ProSA-web Z-score of -3.26 falls within the range characteristic of experimentally determined proteins of comparable length, although it lies toward the lower-density region of the corresponding native-protein distribution. This supports the overall structural plausibility of the model but should be interpreted alongside complementary structural validation metrics rather than as definitive evidence of fold quality [46]. The predicted physicochemical properties suggest a relatively stable and moderately hydrophilic protein, with the aliphatic index providing a theoretical indication of potential thermostability, while divergent solubility scores between DeepSoluE (0.5757) and NetSolP-1.0 (0.8158) reflect known tool variability and underscore the need for empirical solubility testing after expression. Nevertheless, both predictions indicate a favourable likelihood of solubility.

Convergent predictions of antigenicity (VaxiJen v2.0 0.7987, ANTIGENpro 0.7716), non-allergenicity (AllerTOP v2.0, AllergenFP), and non-toxicity (ToxinPred2) constitute the strongest available *in silico* evidence that the deliberate inclusion of *Na*-ASP-2-derived epitopes did not reintroduce the IgE-reactive liability responsible for that antigen’s earlier clinical failure [13]. This should not, however, be equated with demonstrated tolerability; IgE cross-reactivity testing against hookworm-exposed sera remains the definitive assessment.

Population coverage was complete (100%) for MHC class II epitopes but more modest for MHC class I epitopes (63.96% globally; 41.61-66.11% regionally), with South Africa lowest for both classes. This gap most likely reflects the restriction of CTL prediction to the A2, A3, and B7 supertypes rather than a biological shortfall of the epitopes themselves, and argues for broadening the supertype panel or incorporating region-specific HLA frequencies in future iterations, particularly given the disease burden in sub-Saharan Africa.

Protein-protein docking placed the EpiNecaVac-TLR4 complex at a favourable binding free energy (ΔG ≈ -10 kcal/mol; MM/GBSA -63.27 kcal/mol), stabilized by 8 hydrogen bonds and 4 salt bridges across an extensive interface, and normal mode analysis indicated a mechanically coherent rather than loosely associated complex. Together with the built-in adjuvant, this supports a plausible mode of innate immune engagement [37, 38], though it does not by itself establish that TLR4 activation will reach a magnitude sufficient for adjuvant effect *in vivo*; functional cell-based or animal assays are needed to confirm this.

Immune simulation produced an immune-response profile consistent with features commonly associated with effective prime-boost vaccination: an IgM-dominated primary response followed by rising IgG1, IgG2, and combined IgG+IgM titres with declining antigen concentration, expansion of helper and cytotoxic T-cell populations, and evidence of memory formation, alongside sustained IFN-γ and TH signaling consistent with a mixed cellular-humoral profile. These outputs describe the qualitative shape of the response that the construct is designed to elicit and should be treated as hypothesis-generating, pending confirmation in animal models, rather than as a quantitative estimate of protective efficacy.

## Limitations

This work is exclusively computational, and the accuracy of the underlying prediction tools bounds its conclusions. The moderate class I population coverage, the borderline ProSA-web Z-score, and the divergent solubility predictions are points of residual uncertainty rather than settled findings. Mouse MHC class II epitope counts also varied considerably across the seven murine alleles tested (2 to 56 epitopes), which should inform strain selection for any subsequent preclinical immunization study. Recombinant expression, purification, and *in vitro* testing of immunogenicity and IgE cross-reactivity against sera from hookworm-endemic populations are the necessary next steps to substantiate the predictions reported here.

## Conclusion

By integrating computationally filtered epitopes from the three clinical-stage *N. americanus* antigens with a TLR4-agonist adjuvant, EpiNecaVac was designed as a multi-epitope vaccine candidate. The *in silico* analyses predicted favourable properties, including antigenicity, stability, non-allergenicity, non-toxicity, and potential interactions with innate and adaptive immune components. These findings provide a computational basis for further investigation and illustrate how immunoinformatics may support the rational selection and combination of antigenic epitopes, including potentially useful epitopes from previously discontinued vaccine candidates. However, all reported properties and immune-related outcomes remain predictive and have not been experimentally demonstrated. Therefore, experimental studies remains essential before any claims regarding safety or protective efficacy in humans can be made.

## Supporting information

**S1_File**: Table 1. Characteristics of predicted conformational epitopes.

**S1_File**: Figure 1. Population Coverage Analysis for MHC Class I Global and Regional Assessments.

**S1_File**: Figure 2. Population Coverage Analysis for MHC Class II Global and Regional Assessments.

## Acknowledgements

The authors acknowledge all the open-source and globally available databases utilized for data mining.

## Authors’ contributions

**Conceptualization,** Y.A.S.T.; sand R.A.S.; **Data curation,** Y.A.S.T.; R.A.S.; S.M.G.; L.M.S., G.T.N.; T.M.E.; K.P.N.; M.N.M.; and E.N.; **Methodology**, Y.A.S.T.; R.A.S., S.M.G.; **Software**, Y.A.S.T.; R.A.S.; G.T.N., K.P.N.; T.M.E.; J.A.C .N.; and M.N.M.; **Formal analysis**, Y.A.S.T., R.A.S.; L.M.S.; M.N.M.; K.P.N.; G.T.N.; and E.N.; **Investigation,** Y.A.S.T.; R.A.S.; L.M.S.; and K.P.N.; **Resources,** Y.A.S.T.; S.M.G.; and R.A.S.; **Writing - Original Draft Preparation,** Y.A.S.T.; and R.A.S.; **Writing - Review and Editing**, All authors.; **Visualization,** R.A.S.; T.Y.A.S., L.M.S., and **Supervision**, S.M.G., R.A.S.; **Project Administration**, S.M.G.; and R.A.S.; All authors have read and agreed to the published version of the manuscript.

