## Supplementary files for "Immunoinformatics-Based Design of a Chimeric Vaccine Against *Necator americanus* Targeting the *Na*-APR-1, *Na*-GST-1, and *Na*-ASP-2 Vaccine Candidates"

**Table 1. Characteristics of predicted conformational epitopes.**

| No. | Residues | Number of residues | Score |
| --- | --- | --- | --- |
| 1 | A:E427, A:K428, A:G429, A:E430, A:P431, A:C432, A:S433, A:K434 | 8 | 0.976 |
| 2 | A:G446, A:V447, A:G448, A:Q449, A:N450, A:L451, A:K452, A:L453, A:T454 | 9 | 0.957 |
| 3 | A:C435, A:E436, A:N437 | 3 | 0.955 |
| 4 | A:K438, A:G440, A:E441, A:L442, A:A443, A:E444 | 6 | 0.955 |
| 5 | A:V414, A:P415, A:E416 | 3 | 0.864 |
| 6 | A:M410, A:L411, A:T412, A:F413 | 4 | 0.84 |
| 7 | A:E394, A:D395, A:G396, A:R397 | 4 | 0.825 |
| 8 | A:I243, A:L244, A:A245, A:K246, A:A248, A:A249, A:N250, A:G251, A:P252, A:G253, A:P254, A:G255, A:G256, A:G257, A:V258 | 15 | 0.825 |
| 9 | A:E265, A:P266, A:I267, A:T268 | 4 | 0.821 |
| 10 | A:G464, A:E465, A:L466, A:A467, A:E468 | 5 | 0.778 |
| 11 | A:G472, A:Q473, A:K474, A:K475, A:H476, A:H477, A:H478, A:H479, A:H480, A:H481 | 10 | 0.762 |
| 12 | A:M1, A:A2, A:K3, A:L4, A:S5, A:T6, A:D7, A:E8, A:L9, A:L10, A:D11, A:A12, A:F13, A:K14, A:E15, A:M16, A:T17, A:L18, A:L19, A:E20, A:L21, A:S22, A:D23, A:V25, A:K26 | 25 | 0.761 |
| 13 | A:G455, A:K456, A:A458, A:S459, A:K460, A:A461 | 6 | 0.757 |
| 14 | A:D259, A:R261, A:R262 | 3 | 0.745 |
| 15 | A:F24, A:K27, A:F28, A:E29, A:E30, A:T31, A:F32, A:T35 | 8 | 0.728 |
| 16 | A:M399, A:A400, A:I401, A:K402, A:K403, A:S404, A:E405, A:H406, A:N407, A:A408, A:T409 | 11 | 0.714 |
| 17 | A:T270, A:P271, A:T273 | 3 | 0.714 |
| 18 | A:A41, A:V42, A:A43, A:A44, A:A45, A:G46, A:A47, A:A48, A:P49, A:A50 | 10 | 0.642 |
| 19 | A:G360, A:Q361, A:A362, A:K363, A:D364, A:G365, A:K366, A:K367, A:S368, A:G369 | 10 | 0.64 |
| 20 | A:A336, A:L337, A:G338, A:Q339, A:A340, A:K341, A:D342, A:G343, A:G344, A:P345, A:G346, A:P347, A:G348 | 13 | 0.611 |
| 21 | A:A229, A:A230, A:Y231, A:Y232, A:K234, A:Q235 | 6 | 0.604 |
| 22 | A:A218, A:Y219, A:G222, A:K223, A:D224, A:I225, A:E227 | 7 | 0.603 |
| 23 | A:Y237, A:H238, A:K241 | 3 | 0.599 |
| 24 | A:A70, A:G71, A:D72, A:K73, A:K74, A:G76, A:V77, A:I78, A:K79, A:V80, A:R82, A:E83, A:V85, A:S86, A:G87, A:L88, A:G89, A:L90, A:A93, A:K94, A:V97, A:D98, A:G99, A:K119, A:A122, A:A123, A:G124, A:A125 | 28 | 0.576 |

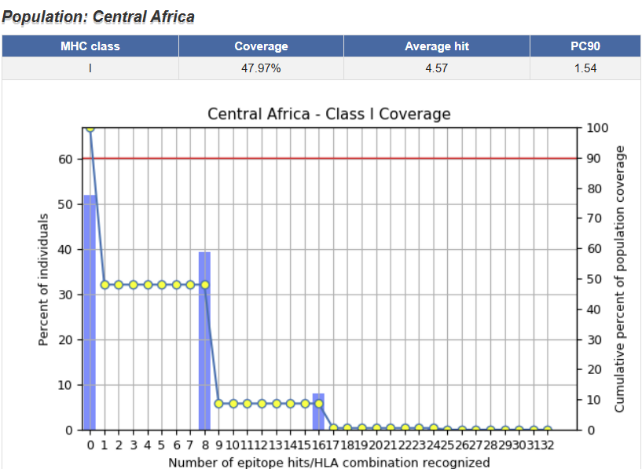

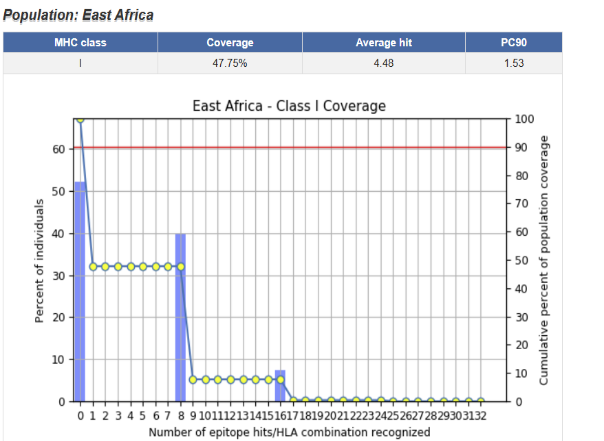

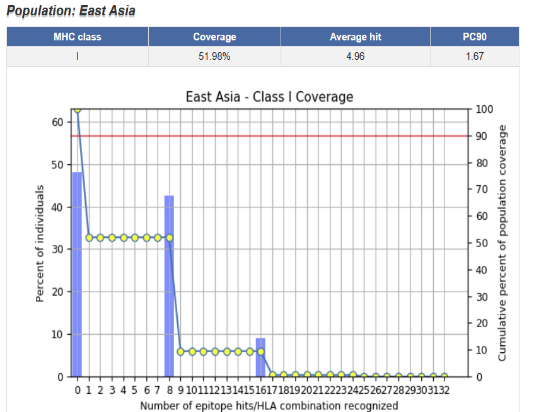

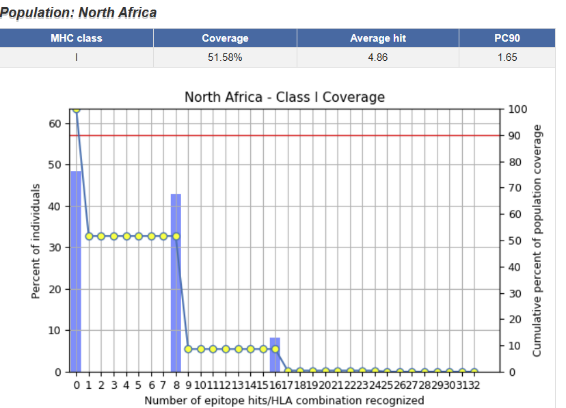

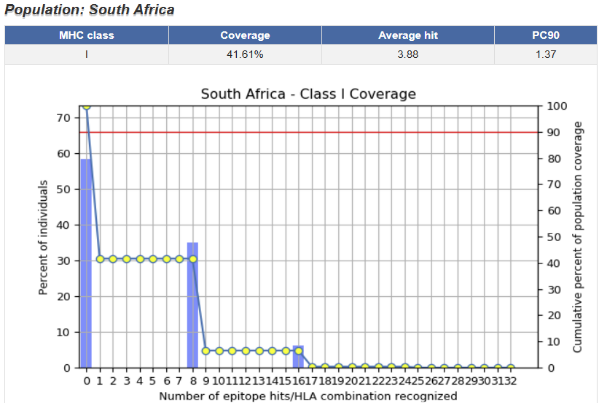

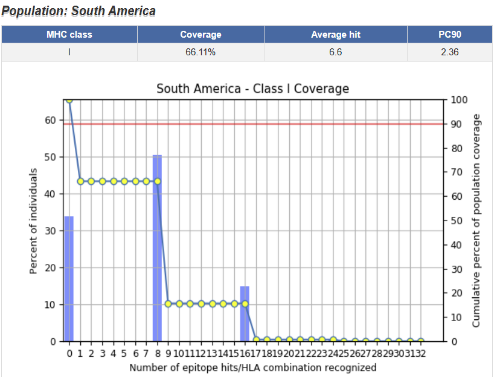

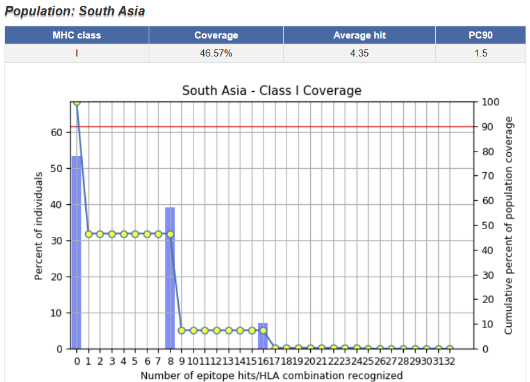

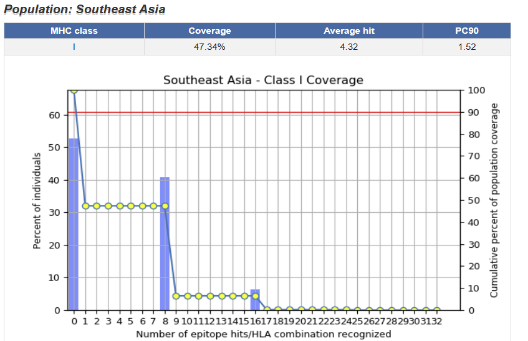

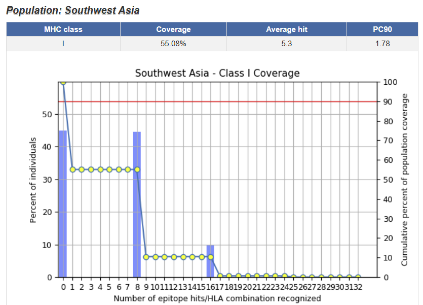

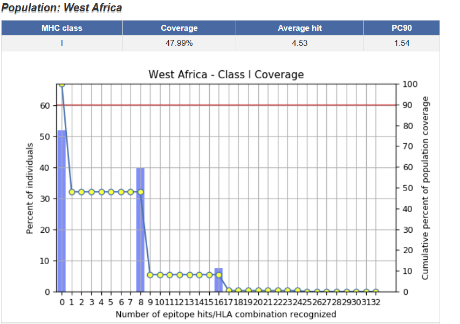

***Figure 1.*** ***Population coverage analysis for MHC class I global and regional*** ***assessments.*** *Population coverage analysis of predicted T-cell epitopes across the global population is presented as follows: MHC class I epitope coverage demonstrates a global population coverage of 63.96%, with average hit and PC90 values suggesting extensive HLA representation. The histograms and cumulative curves illustrate the percentage of individuals recognizing a specific number of epitope-HLA combinations.*

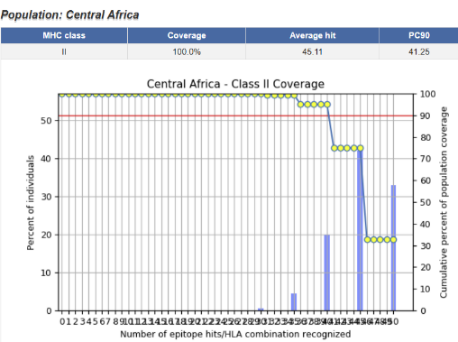

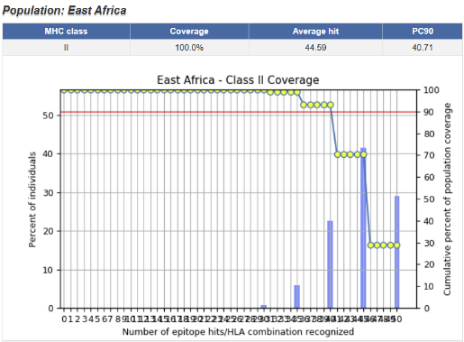

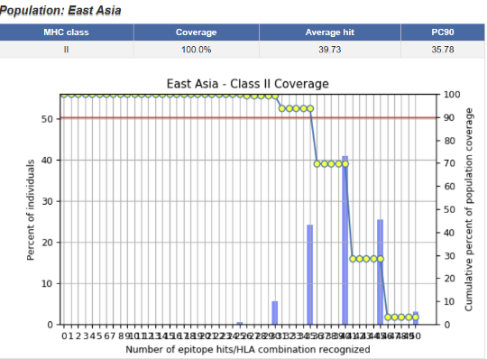

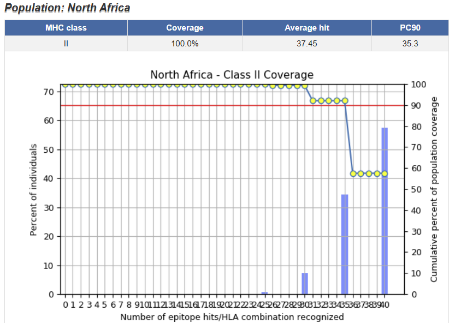

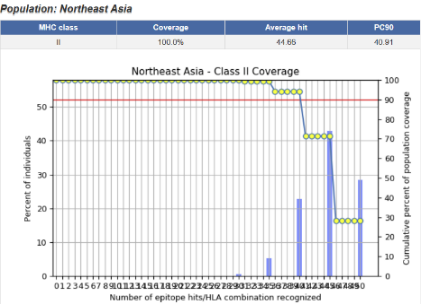

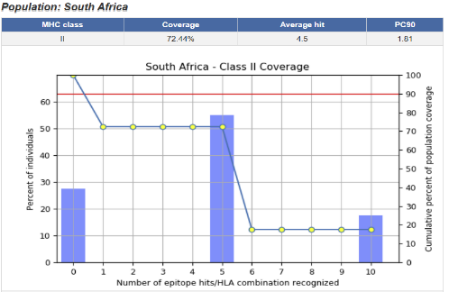

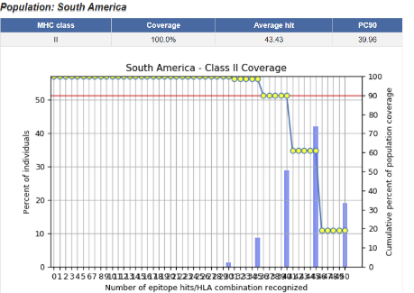

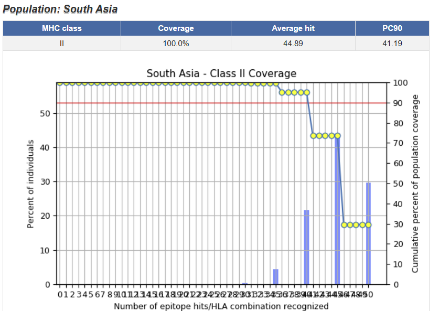

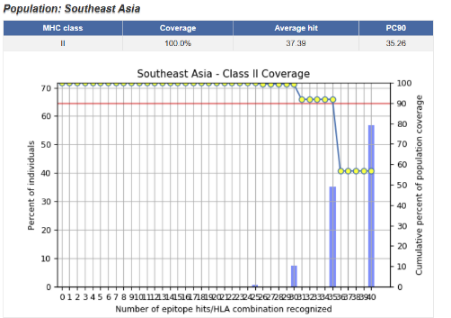

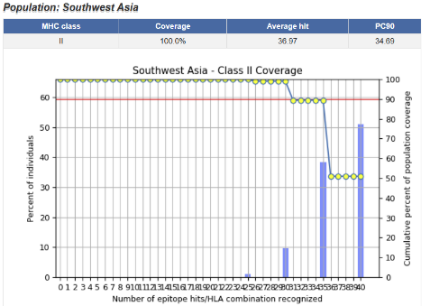

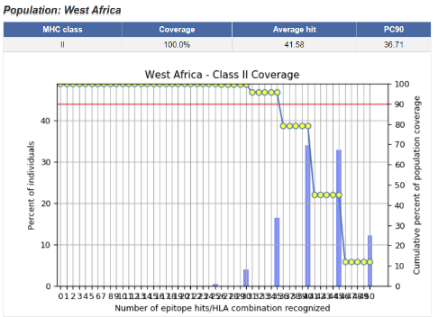

***Figure 2****.* ***Population coverage analysis for MHC class II global and regional*** ***assessments.*** *Population coverage analysis of predicted T-cell epitopes across the global population is presented as follows: MHC class II epitope coverage reflects a global population coverage of 100%, highlighting the distribution of HLA class II alleles across various populations. The histograms and cumulative curves illustrate the percentage of individuals recognizing a specific number of epitopes-HLA combinations.*
